# Lipids intercalate and mediate multi-channel assemblies of connexin-46/50 gap junctions

**DOI:** 10.64898/2026.05.28.728492

**Authors:** Connor S. Garrels, Janette B. Myers, Samson A. Souza, Joshua M. Jarodsky, Steve L. Reichow

## Abstract

Gap junction channels enable direct electrical and metabolic exchange between adjacent cells and tissues, where they organize into dense plaques containing tens to thousands of channels. Although plaque formation is known to modulate junctional conductance, the structural basis for channel–channel organization within a membrane environment remains poorly defined. Here, we reconstitute native lens connexin-46/50 (Cx46/50) gap junction channels into MSP-based lipid nanodiscs that incorporate multiple channels to create miniature plaque-like complexes suitable for single-particle cryo-electron microscopy (cryo-EM). We determine high-resolution structures of dual-channel assemblies in two distinct configurations and find no ordered protein–protein contacts across the interface. Instead, the inter-channel space is occupied by ordered lipid density, indicating that channel packing in these assemblies is lipid mediated. These channel–channel interfaces stabilize discrete lipid populations, including an interstitial lipid intercalated between subunits and positioned near the N-terminal gating domain, suggesting a route by which channel organization could promote lipid occupancy near the pore. Additionally, we leverage this dataset to refine the Cx46/50 single-channel structure to 1.8 Å resolution, revealing exceptional chemical detail of the pore-lining landscape in the stabilized open-state. Together, these results define principles of lipid-mediated multi-channel organization and suggest how plaque-like packing may tune gap junction function through specific lipid interactions.

## INTRODUCTION

Connexin gap junction channels directly couple the cytoplasm of adjacent cells, enabling electrical and metabolic exchange across tissues^1,2^. *In vivo*, gap junction channels rarely operate in isolation. Instead, they coalesce into densely packed membrane arrays known as plaques, containing tens to thousands of channels. This mesoscale organization parallelizes gap junction function and increases the capacity and efficiency of intercellular communication. Plaque formation is therefore a defining feature of gap junction biology, but the structural interactions that govern channel organization within membranes and how this organization influences channel function remain poorly understood. Defining these principles is fundamental for interpreting how connexin dysfunction contributes to diverse human diseases, including deafness, blindness, skin disorders, neuropathies, cancer, and arrhythmia^3–6^.

Gap junction intercellular channels form by the docking of two hexameric hemichannels (connexons), yielding a dodecameric complex in which each connexin subunit contributes four transmembrane helices (TM1–4) and two extracellular docking domains (EC1–2) that bridge the intercellular gap^7,8^. Among the most functionally important regions is the cytosolic N-terminal (NT) domain, which folds into the pore vestibule and is a central determinant of channel selectivity and gating^9–14^. Recent high-resolution structures of individual connexin channels obtained by X-ray crystallography and single-particle cryo-electron microscopy (cryo-EM) have provided a detailed basis for understanding pore architecture and conformational transitions associated with the NT^8,15–18^ (reviewed^19^). Our previous work on connexin-46/50 (Cx46/50) showed that in the conductive open-state, the NT adopts a well-ordered α-helical configuration stabilized by hydrophobic interactions with the pore lumen, supporting a wide pore (∼1.4 nm diameter) compatible with permeation of ions and small metabolites^15,16^.

Consistent with these structural observations, single-channel electrophysiology of gap junction channels at minimal transjunctional voltage (V*_j_*) report high open-state probabilities^20,21^. In contrast, elevated V*_j_* and diverse cytoplasmic signals can promote gating transitions that reduce junctional conductance^22–24^ (reviewed^25^). Recent structural studies on Cx46/50 conducted under gating conditions, such as acidic pH or elevated calcium, show the NT can become destabilized and adopt conformations expected to restrict or occlude the pore^17,26^. Notably, lipids can also occupy or stabilize conformations of the NT associated with gating in Cx46/50, suggesting a plausible route by which the membrane environment may directly influence channel state^17^.

Functional observations in tissues point to an additional layer of regulation that is difficult to reconcile with observations of isolated channels. Across multiple studies in lens, cardiac, and neuronal tissues, as well as in model cell systems, plaque conductance is often substantially lower than predicted based on plaque size and the unitary conductance of open channels, implying that many channels within plaques are closed or non-conductive under basal conditions^27–33^. Strikingly, even relatively small plaques have been proposed to contain few, if any, functional channels despite harboring large numbers of connexins^30^. These results suggest that plaque organization may tune gap junction function, but the molecular basis for this regulation, including the possible roles of channel packing and local lipid organization, remains unclear.

Early electron microscopy and 2D-crystallography studies established that gap junction plaques can form ordered or quasi-ordered hexagonal lattices, exhibiting varying degrees of local heterogeneity in channel spacing and organization^34–39^. These approaches provided foundational measurements of intermembrane separation and lattice parameters and were later extended to 2D protein–lipid crystals, yielding low- to intermediate-resolution views of channel architecture in membrane environments^7,18,40,41^. However, the limited resolution and ensemble averaging inherent to these methods precluded identification of the specific interactions that govern channel–channel packing, leaving questions as to whether lipids contribute directly to stabilizing distinct packing modes or modulating pore features linked to gating unresolved.

Here, we develop a nanodisc reconstitution strategy that incorporates multiple native lens Cx46/50 intercellular channels into MSP-based lipid nanodiscs compatible with high-resolution single-particle cryo-EM. The resulting particles frequently contain two full dodecameric channels within shared nanodiscs and also sample rare higher-order assemblies, enabling direct interrogation of channel–channel organization in a defined lipid environment. We determine high-resolution structures of dual-channel assemblies in two distinct packing configurations and find that channel association occurs through a lipid-occupied interface rather than ordered protein–protein contacts. These interfaces stabilize multiple lipid populations, including an interstitial lipid intercalated between subunits and positioned near the NT gating region, suggesting a route by which plaque-like packing could influence lipid occupancy near the pore. Further utilizing the same dataset, we also refine a single-channel Cx46/50 structure to 1.8 Å resolution in the stable open-state conformation, revealing unprecedented channel detail and enabling direct structural comparisons between single and dual-channel assemblies. Together, these data define the role of lipids in mediating high-order gap junction assembly and provide a framework for how plaque-like packing may tune gap junction function through specific lipid interactions.

## RESULTS

### Sample preparation: capturing multi-channel Cx46/50 assemblies

Native Cx46/50 gap junction channels were purified from the nuclear region of fresh bovine lenses and reconstituted into MSP1E1 lipid nanodiscs containing dimyristoyl-phosphatidylcholine (DMPC) (**Fig. 1a,b**). Under reconstitution conditions similar to those we have previously used to generate predominantly single-channel particles^15–17,26^, we observed that a subset of assemblies contained more than one intercellular channel within a shared membrane scaffold. To enrich these higher-order assemblies for structural analysis, we collected a broader region of the size-exclusion chromatography (SEC) trace than typically selected for isolating single-channels (**Fig. 1a**).

**Figure 1.**
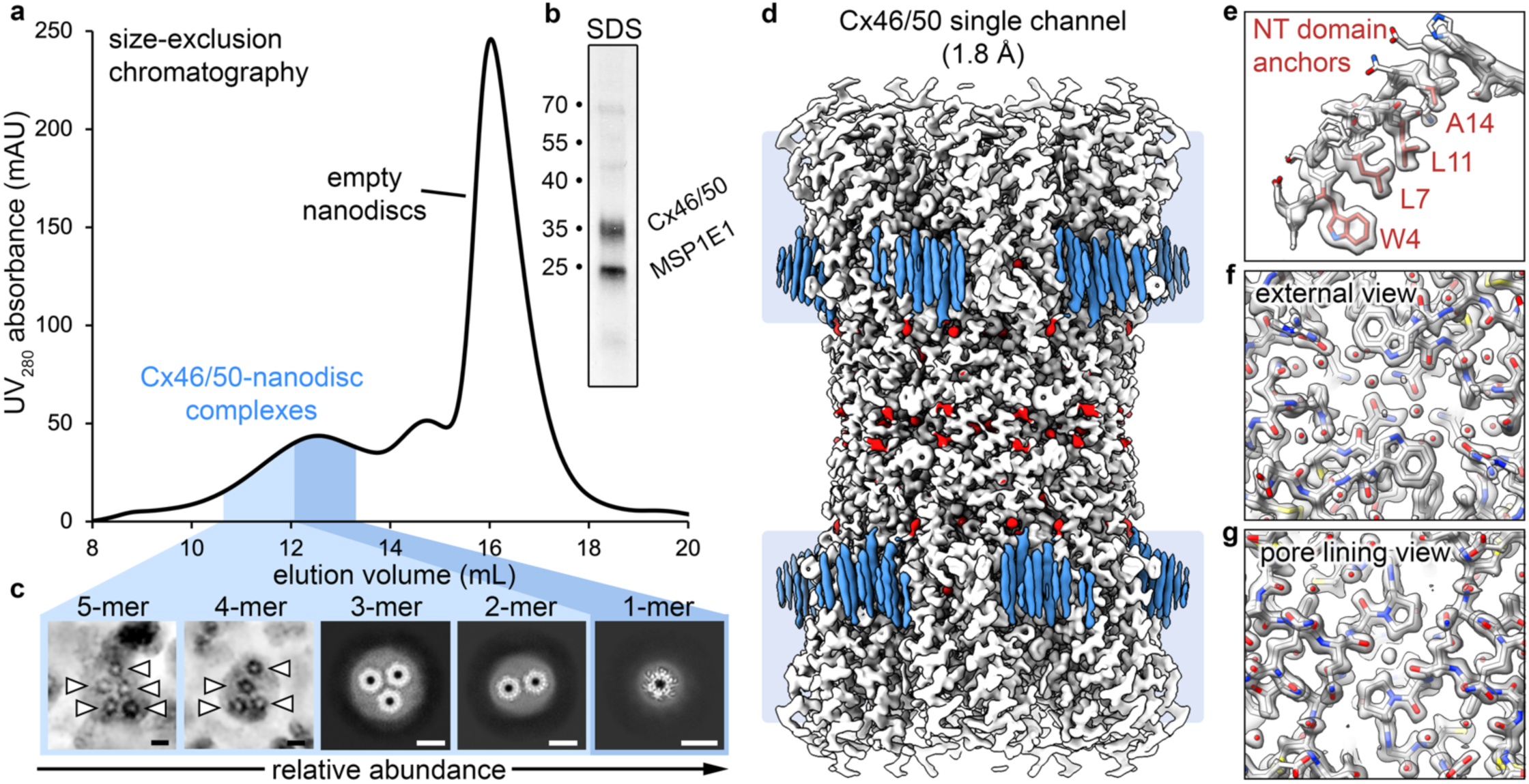
Enrichment of multi-channel Cx46/50 nanodisc assemblies and 1.8 Å single-channel reconstruction. **a,** Size-exclusion chromatography (SEC) trace of the Cx46/50–MSP1E1/DMPC reconstitution. A narrow fraction (dark blue) is enriched for single-channel particles, whereas a broader fraction (light and dark blue regions), pooled for this study, contains single-channel particles together with a heterogeneous mixture of multi-channel assemblies. The major late-eluting peak corresponds to empty nanodiscs. **b,** Silver-stained SDS–PAGE of the reconstitution showing Cx46/50 and MSP1E1. **c,** Representative cryo-EM views of multi-channel assemblies observed in the pooled SEC fraction. Five- and four-channel particles are shown as denoised micrographs (arrowheads indicate individual channels), whereas three-, two-, and single-channel particles are shown as representative 2D class averages. The arrow indicates relative abundance across observed assembly sizes. Scale bars, 10 nm. **d,** Cryo-EM reconstruction of the single-channel Cx46/50 assembly at 1.8 Å resolution. Density corresponding to the protein (white), annular lipids (blue), and ordered solvent features (red). Light blue boxes indicate the approximate membrane boundaries. **e-g,** Representative map-to-model views illustrating reconstruction quality. *(e)* NT domain in the stabilized open state with hydrophobic anchor residues labeled (red). *(f)* Extracellular (external) view at the intercellular docking interface. *(g)* Pore-lining view from within the channel. Models in *(e–g)* are colored by atom type.

Preliminary single-particle cryo-EM analysis of the broadened SEC fraction revealed a heterogeneous population containing both single-channel particles and multi-channel assemblies in which two or more dodecameric channels are incorporated within a shared lipid bilayer (**Fig. 1c; Supplementary Fig. 1**). Although canonical MSP1E1 nanodiscs are typically ∼10–12 nm in diameter^42^, we also observed substantially larger membrane particles, in some cases exceeding ∼50 nm. Two- and three-channel assemblies were sufficiently abundant to be readily identified in 2D class averages and initial 3D reconstructions, whereas larger assemblies containing four or more channels were observed in raw micrographs but were comparatively rare (**Fig. 1c; Supplementary Fig. 1**). We interpret these as MSP-stabilized membrane particles that likely incorporate additional MSP scaffold proteins, enabling accommodation of multiple channels within a shared bilayer. We also occasionally observed assemblies with three membranes, consistent with non-physiological ‘head-to-tail’ channel arrangements, similar to what has been reported in prior 2D electron crystallography studies^18^ (**Supplementary Fig. 1**).

Although single-channel particles remained the dominant species overall (∼75% of classified particles), dual-channel assemblies were the most abundant multi-channel species (∼25% of classified particles) and were therefore carried forward for high-resolution refinement. Three-channel assemblies could be reconstructed but remained limited by particle number (<1% of classified particles**; Supplementary Fig. 1**). In the following sections, we compare the high-resolution single-channel structure obtained from this dataset with the dual-channel assemblies to define the features that emerge in a plaque-like packing context.

### Cx46/50 single-channel assembly at 1.8 Å resolution

To provide an internal reference for comparative analysis to the dual-channel assemblies, we separately processed the abundant single-channel particles present in the same dataset (**Fig. 1d–g; Supplementary Figs. 2,3; Supplementary Table 1**). This reconstruction refined to 1.8 Å global resolution (gold-standard FSC), improving upon our previously reported 1.9 Å ovine Cx46/50 structure obtained under comparable non-gating conditions^16^. Atomic models for both Cx46 and Cx50 were refined into the cryo-EM map. As in prior work^15,16^, the heteromeric/heterotypic arrangement of Cx46 and Cx50 is not distinguishable in the density despite various attempts of classification and alternative symmetric and asymmetric refinements, reflecting the >80% sequence identity across structured regions of the two isoforms and limitations of current cryo-EM classification methods (**Supplementary Fig. 3**). Due to this high sequence and structural homology, we use Cx46 as the archetype model for the following analyses unless stated otherwise.

Comparison with our prior open-state Cx46/50 models^16^ showed no substantial differences in overall architecture, including the transmembrane helix arrangement and the stabilized open-state NT conformation (Cα r.m.s.d. = 0.2 Å; **Supplementary Fig. 3**), consistent with the close sequence identity between these species (∼97% identical over the modeled regions). Annular lipid features belonging the extracellular leaflet also display similar long-range stabilization relative to the protein surface, providing a consistent baseline for comparisons to dual-channel assemblies (**Fig. 1d**; **Supplementary Fig. 3**).

While the overall architecture of Cx46/50 is consistent with our previous reports, the improved resolution achieved here provides markedly enhanced chemical detail, enabling confident interpretation of pore-lining stereochemistry and local interactions that shape conductance and permeation (**Supplementary Movie 1**). Sidechain rotamers at the NT anchoring site and along the pore-lining surface are more clearly defined, and coordinated networks with ordered water molecules are apparent within the channel lumen and at the extracellular junctional interface (**Fig. 1e-g**). Together, these features provide a high-confidence structural baseline for interpreting potential differences in lipid occupancy and pore-lining chemistry in dual-channel assemblies.

### Resolving dual-channel Cx46/50 assemblies from a heterogenous mixture

Initial template-based particle picking using a preliminary dual-channel map as a reference yielded substantial contamination by single-channel particles, as evident from 2D classification and 3D reconstructions in which one channel was consistently better resolved than the other (**Supplementary Fig. 4**). To separate bona fide dual-channel particles from single-channel contaminants, we implemented a combination of heterogeneous refinement and multiple rounds of 3D variability analysis (3DVA) as particle curation steps (**Fig. 2a**).

**Figure 2.**
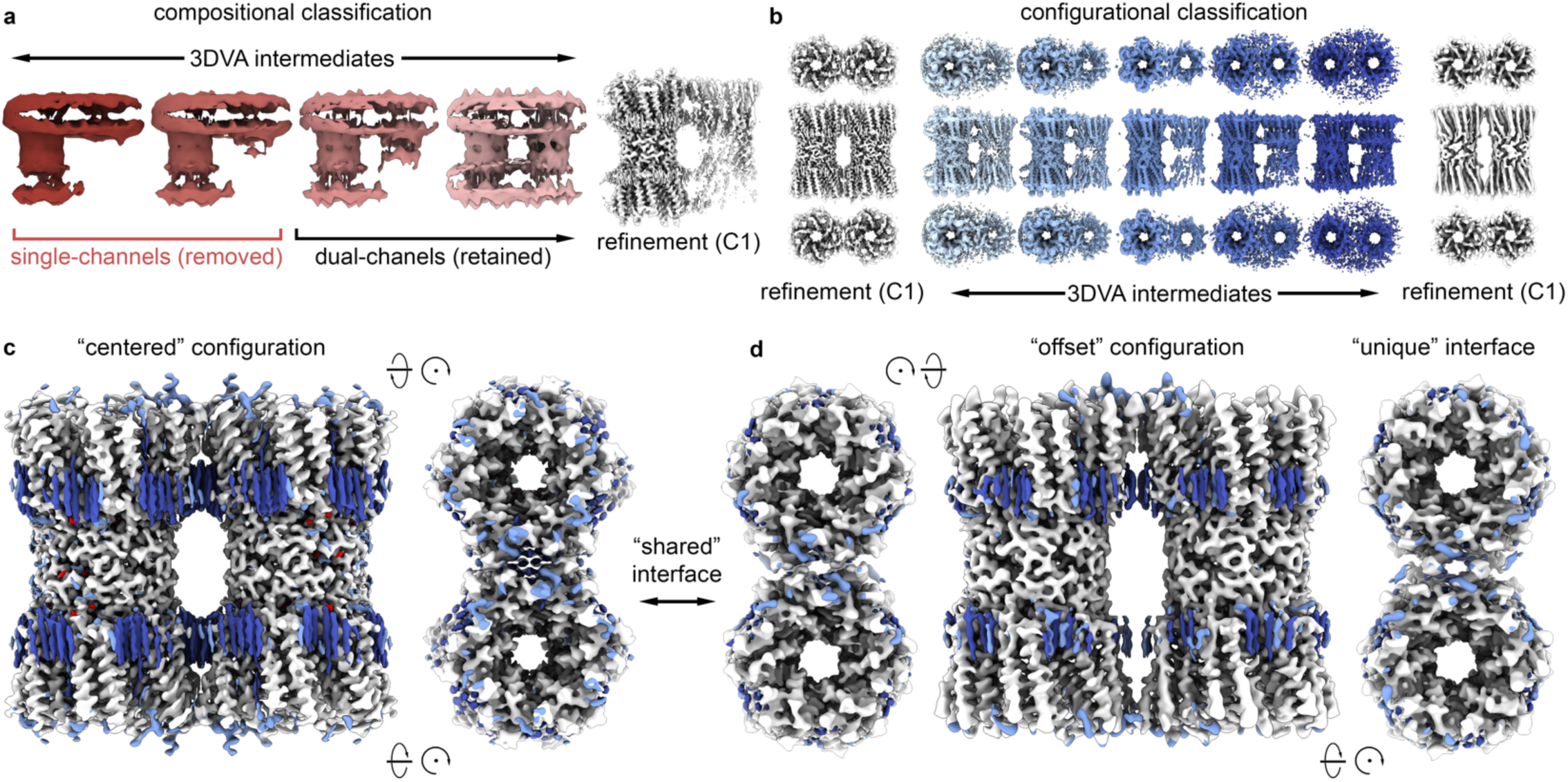
Classification of dual-channel Cx46/50 assemblies reveals two packing configurations. **a,** Schematic of compositional separation by 3DVA. Four representative intermediates along the dominant principal component are shown (red), spanning a continuum from reconstructions dominated by single-channel particles (left) to those dominated by dual-channel particles (right). The single-channel end of the component was removed, and the dual-channel-enriched subset was retained for refinement. C1 consensus refinement following 3DVA yields an asymmetric reconstruction (white) in which one channel is better resolved than the other, indicating additional heterogeneity beyond composition. **b,** Schematic of configurational heterogeneity separated by 3DVA. Five representative intermediates along the dominant principal component are shown (blue), spanning two endpoint configurations. Particles corresponding to each end of the component were refined independently without applied symmetry (white). **c-d,** Final cryo-EM maps of dual-channel assemblies in the centered configuration *(c)* refined with D2 symmetry and offset configuration *(d)* refined with C2 symmetry. The centered assembly contains equivalent (“shared”) channel–channel interfaces at both cytosolic faces, whereas the offset assembly contains one “shared” interface and one “unique” interface (rightmost panel). Maps are shown with protein density in white; ordered lipid features in dark blue; ordered solvent features in red; and additional unassigned density in light blue.

To perform an initial coarse separation of single- and dual-channel particles, we carried out one round of ‘baited’ heterogeneous refinement (**Supplementary Fig. 4**). In this scheme, the heterogeneous particle stack was classified against competing dual-channel and single-channel reference volumes. The particles assigned to the single-channel class were discarded and those assigned to the dual-channel class were retained. This approach effectively removed most single-channel contaminants, but residual contamination persisted despite additional rounds. We therefore leveraged 3DVA as the primary downstream curation strategy.

The dominant principal component from the first round of 3DVA robustly separated remaining single-channel particles from dual-channel particles (**Fig. 2a, Supplementary Fig. 4, Supplementary Movie 2**). The half of the component containing dual-channel particles was kept and the other half discarded. Assessment by 2D classification confirmed that no detectable single-channel particles remained. However, a consensus refinement of the dual-channel particles still contained only one completely resolved channel, indicating persistent heterogeneity beyond single-channel contamination (**Fig. 2a, Supplementary Fig. 4**).

A second round of 3DVA revealed two distinct configurations of the dual-channel assembly were responsible for the observed heterogeneity (**Fig. 2b, Supplementary Fig. 4, Supplementary Movie 3**). Particles belonging to each half of the 3DVA principal component were isolated and refined separately, yielding two distinct dual-channel reconstructions. The reconstructions each contained two fully resolved channels, indicating all remaining heterogeneity had been accounted for, and were carried forward to high-resolution refinement (**Fig. 2b)**.

One dual-channel orientation, comprising ∼55% of the dual-channel population, refined with D2 symmetry to a global resolution of 2.9 Å (**Fig. 2c; Supplementary Fig. 5; Supplementary Table 1**). In this arrangement, the channels adopt a small relative tilt in which the relative tilt axis passes through the center of the extracellular docking interface, resulting in equivalent cytosolic-facing interfaces on both sides of the paired assembly. We refer to this configuration as the ‘centered’ assembly. The centered map resolves density for the two channels as well as associated DMPC lipids, predominantly acyl chains with partially or fully resolved headgroups in regions of higher local resolution (**Fig. 2c**). Density consistent with ordered solvent is also apparent, with solvent features most prominent in regions of higher local resolution away from the channel–channel interface (**Supplementary Fig. 5**).

The second orientation refined with C2 symmetry to a global resolution of 3.4 Å (**Fig. 2d; Supplementary Fig. 5; Supplementary Table 1**). In this arrangement, the channels also adopt a relative tilt, but the relative tilt axis is offset from the central plane of the channel, such that the two cytosolic-facing interfaces are not equivalent. We refer to this conformation as the ‘offset’ assembly. One cytosolic interface closely resembles that observed in the centered assembly (the shared interface), whereas the opposite interface adopts a distinct configuration (the unique interface) (**Fig. 2d**). Comparisons of symmetrized and non-symmetrized (C1) reconstructions confirmed that the differences between the centered and offset assemblies do not arise from symmetry application (**Supplementary Fig. 5**). In the offset map, lipid and solvent features are present but less extensively resolved, consistent with the relatively lower global and local resolution (**Supplementary Fig. 5**).

### Dual-channel assemblies adopt the stabilized open-state conformation

Atomic models for both Cx46 and Cx50 were refined into the centered and offset cryo-EM reconstructions (**Supplementary Fig. 5; Supplementary Table 1**). As in the high-resolution single-channel reconstruction, the heteromeric/heterotypic arrangement of Cx46 and Cx50 could not be distinguished from the density maps, and as such Cx46 is again presented as the archetype model with isoform-specific features highlighted when relevant.

Both the centered and offset assemblies adopt the stabilized open-state, as indicated by the conformation of the NT helix, consistent with prior Cx46/50 structures under non-gating conditions^15,16^ (**Fig. 3a–c**). Alignment of the centered assembly with the 1.8 Å open-state model refined from the same dataset shows that the dual-channel arrangement does not meaningfully alter the NT helix or overall channel architecture (Cα r.m.s.d. = ∼0.3 Å; **Fig. 3b,d; Supplementary Fig. 6**). In particular, the conserved hydrophobic NT anchor residues (W4, L7, L10 (I10 in Cx50), L11 and A14 (V14 in Cx50)) occupy the characteristic packing interactions with TM1/2 that stabilized the NT in the conductive open-state^15,16^ (**Fig. 3d**).

**Figure 3.**
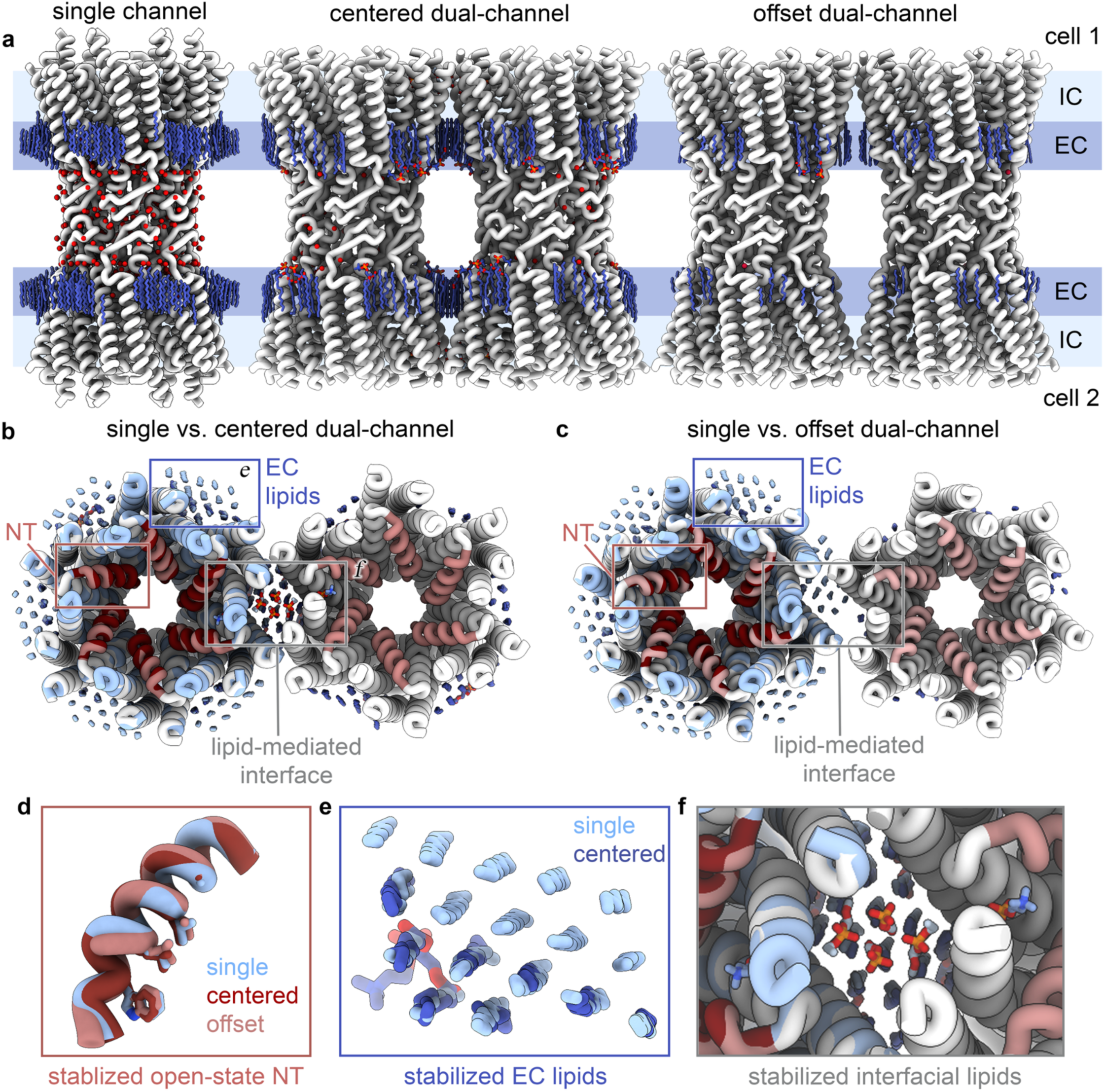
Dual-channel assemblies retain the stabilized open-state and stabilize interface-associated lipids. **a,** Overview of the single-channel, centered dual-channel, and offset dual-channel models. Protein is shown in white; annular lipids in dark blue; and ordered waters in red. Shaded bands denote the approximate membrane boundaries and are labeled for the extracellular-facing (EC) and intercellular-facing (IC) leaflets**. b-c,** Structural alignment of the single-channel Cx46 model (blue and dark red NT) with the centered *(b)* and offset *(c)* dual-channel models (white and light red NT). Boxes indicate regions highlighted in subsequent panels: the NT domain region (red box), conserved extracellular leaflet lipid features (blue box), and the lipid-mediated channel–channel interface (gray box). **d,** Superposition of NT domains from the single-channel (blue), centered (dark red), and offset (light red) assemblies, showing the NT domains all adopt the same stabilized open-state with preserved packing of hydrophobic anchor residues. **e,** Superposition of stabilized extracellular leaflet (EC) annular lipids in the single-channel (light blue) and centered dual-channel (dark blue) assemblies. **f,** Close-up view of the lipid-mediated interface in the centered dual-channel assembly (gray box in *b*), highlighting stabilized interfacial lipids.

Annular DMPC densities, within the extracellular (EC) leaflet, are consistently observed in the single-channel and in both dual-channel assemblies, with acyl-chain positions and long-range hexagonal packing largely conserved across reconstructions (**Fig. 3b,c,e**). In the dual-channel maps, these EC lipids are resolved over a shorter radial extent from the protein surface, consistent with differences in local resolution and/or potential differences in lipid stabilization relative to the 1.8 Å single-channel reconstruction.

The channels in the offset assembly also show similarity in overall architecture to the single-channel conformation (Cα r.m.s.d. = ∼0.4 Å; **Fig. 3c,d; Supplementary Fig. 6**). Importantly, the NT is preserved in the stabilized open-state conformation at both the shared and unique interfaces, indicating that the offset assembly reflects a difference in global channel orientation rather than localized deformation at one face.

Notably, in addition to EC annular lipids, the centered map contains density consistent with lipids at the intracellular (IC) leaflet and an interstitial lipid (IL) intercalated between subunits near the cytoplasmic vestibule (**Fig. 3b,f**). Lipids at these locations were not resolved in our prior single-channel reconstructions of lens Cx46/50 under similar non-gating conditions, or in the single-channel assembly resolved in this study (**Supplementary Fig. 7**), despite its higher global resolution. Although these additional lipids were also not resolved in the offset dual-channel map, this may be explained by the lower overall resolution and reduced local map quality. We describe these uniquely resolved lipid features in detail in the following section.

### Channel association is lipid-mediated and organizes ordered bilayer features

In the dual-channel reconstructions, we do not observe ordered protein–protein contacts across the interface; instead, the inter-channel space is occupied by ordered lipid density (**Fig. 4a–c**). Lipid ordering is most extensive in the centered dual-channel assembly and localizes to the channel–channel interface (**Fig. 4a**).

**Figure 4.**
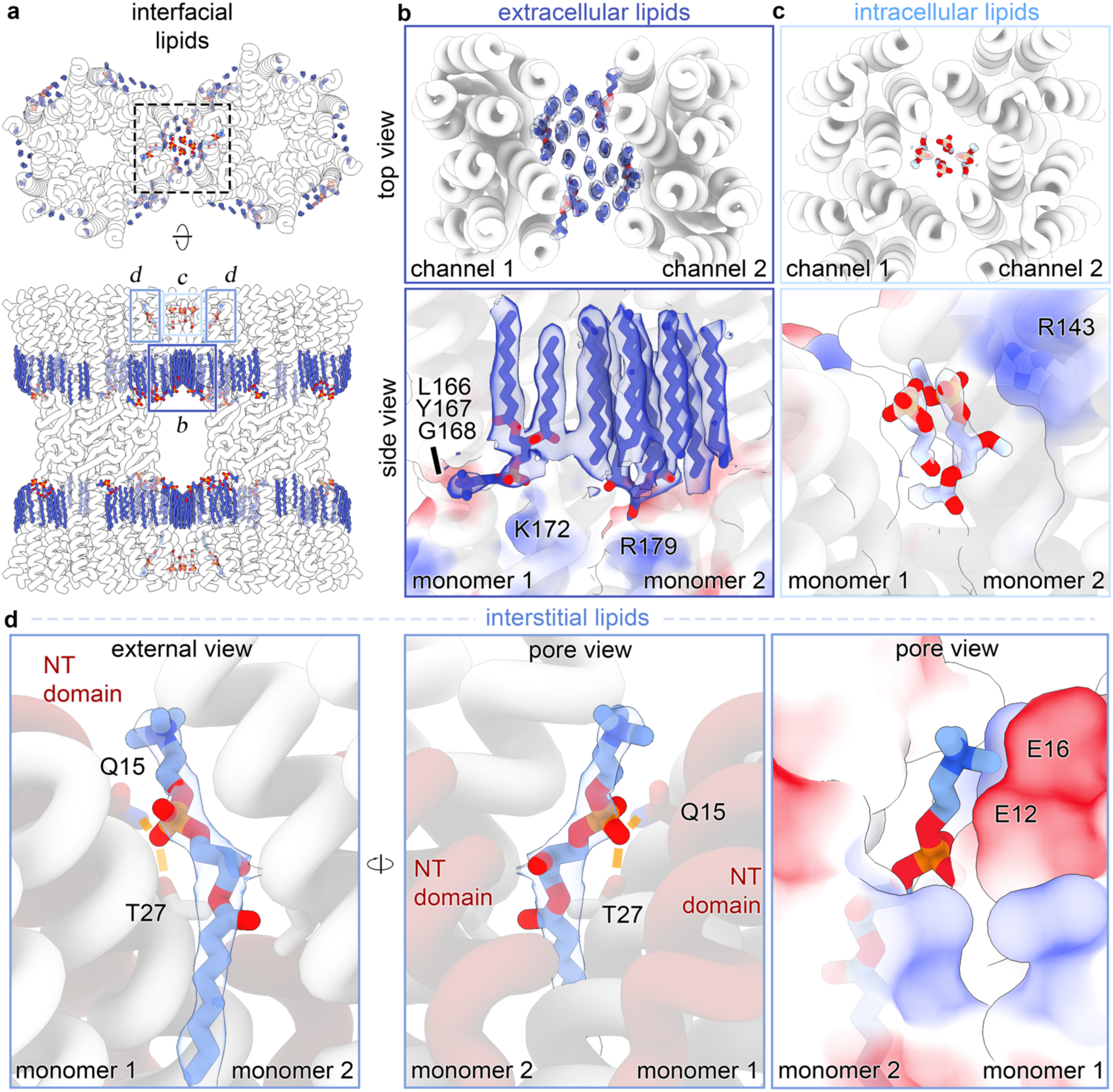
Centered dual-channel assembly stabilizes discrete lipid populations at the lipid-mediated interface. **a,** Overview of lipid features resolved in the centered dual-channel assembly. The lipid-mediated interface stabilizes three lipid populations: extracellular leaflet (EC) lipids (dark blue), intercellular leaflet (IC) lipids (light blue), and an interstitial lipid (IL) at the interface (medium blue). Boxed regions indicate the locations of zoom views shown in b–d. **b,** Zoom-views of the stabilized EC leaflet lipids that occupy the full inter-channel space at the interface. Top (upper) and side (lower) views show ordered acyl-chain density spanning between channels, with partial or complete headgroup density visible at selected sites. Modeled headgroup features are positioned near charged surface patches, including an electronegative backbone region (L166–Y167–G168) and nearby basic residues (K172 and R179) indicated by semi-transparent surface potential in the lower panel. **c,** Zoom-views of the IC leaflet lipid headgroup features at the interface. Four partial headgroup densities are resolved in the IC leaflet at the centered interface and are not apparent in the single-channel reconstruction. Two of these features are positioned adjacent to R143 (shown), consistent with electrostatic stabilization of the phosphate region. **d,** Zoom-views of the IL lipid at the centered interface, intercalated between neighboring subunits near the cytoplasmic vestibule. The phosphocholine headgroup projects toward the pore vestibule, while a short segment of one acyl chain extends into the IC leaflet region. The IL headgroup forms polar contacts with Q15 on the NT (N15 in Cx50) and T27 on TM1, and is positioned adjacent to an electronegative patch contributed by E12 and E16.

In the EC leaflet, each connexin monomer is associated with an average of ∼8 rod-like densities modeled as DMPC acyl chains, truncated to match the cryo-EM density (**Fig. 4a,b; Supplementary Fig. 7**). In several cases, paired acyl chains are connected by additional density consistent with partial or complete DMPC headgroups (**Fig. 4b**). Headgroup density attributed to the choline moiety lies near an electronegative protein surface patch formed by the backbone carbonyls of L166, Y167, and G168 (L179, Y180, G181 in Cx50), whereas the phosphate region is positioned adjacent to positively charged residues, K172 and R179 (equivalent to L185 and R192 in Cx50) (**Fig. 4b**). All residues are conserved between isoforms except K172/L185. In several cases, the headgroup orientations closely match headgroup-containing lipid classes from previous work^16^ (**Supplementary Figure 7**). Notably, EC leaflet lipids occupy the full space between neighboring channels, adopting a continuous hexagonal packing arrangement (**Fig. 4b**). In the offset dual-channel assembly, EC leaflet lipid headgroup features are also present but less well-defined, again, consistent with the lower local map quality in this reconstruction (**Supplementary Fig. 5**).

In addition to EC leaflet lipids, four oblique densities are uniquely resolved at the IC leaflet of the channel–channel interface, interpreted as IC lipid headgroups (**Fig. 4c**). Based on the z-height and proximity to the guanidinium group of R143 (R156 in Cx50), these densities were considered most consistent with modeling the phosphate region of a DMPC headgroup at these sites. The spacing of these IC leaflet densities mirrors that of the EC leaflet lipid features across the same interface (**Fig. 4c**), consistent with an environment stabilized by channel–channel proximity that locally orders the headgroups of these interfacial lipids.

### An interstitial lipid is stabilized near the NT gating domain

An additional interstitial lipid (IL) is resolved in the centered dual-channel assembly, intercalated between neighboring subunits near the cytoplasmic vestibule proximal to the NT gating domain (**Fig. 4a,d**). The density accommodates a complete phosphocholine headgroup and an approximately seven-carbon segment of one acyl chain, consistent with DMPC (**Fig. 4d**). The headgroup projects toward the pore vestibule at the interface between TM1 of one monomer and TM2 of the adjacent monomer. In the refined model, the lipid headgroup forms polar interactions with Q15 on the NT helix (N15 in Cx50) and T27 on TM1, while the choline moiety lies adjacent to an electronegative region contributed by E12 and E16 on the NT helix (**Fig. 4d**).

Q15 is positioned to contact the IL headgroup only when the NT helix adopts the stabilized open-state conformation. Supporting this assertion, alignment with the pH-gated Cx46/50 structure shows that the gated NT helix would sterically clash with the IL headgroup density in the repositioned gated conformation^17^. Interstitial lipids reported in other gap junction structures occupy positions distinct from the IL site observed here^43–47^ (**Supplementary Fig. 7**), suggesting a unique chemical environment required for stabilization in this state. Although density consistent with lipid occupancy is present at the IL site across multiple subunit interfaces in the centered reconstruction, only the site positioned at the channel–channel interface sufficiently resolved to support confident modeling of the headgroup (**Supplementary Fig. 7**).

Notably, when the IL site is examined in the 1.8 Å single-channel reconstruction, very weak density can be observed that overlaps with the acyl-chain region of the IL site, consistent with partial occupancy, but no density corresponding to the IL headgroup is observed (**Supplementary Fig. 7**). Together, these comparisons support the conclusion that the IL headgroup is uniquely stabilized in the plaque-like packing context captured by the centered dual-channel interface, establishing a structural link between multi-channel organization and lipid occupancy near the NT gating domain.

### Two interface geometries with distinct packing consequences

The centered and offset dual-channel assemblies represent two possibilities for the relative orientations of gap junction channels in a lipid membrane and, as a result, confer two distinct channel–channel interface geometries. In the centered assembly, the two cytosolic-facing interfaces are equivalent (two shared interfaces), whereas in the offset assembly one cytosolic interface matches the centered interface (the shared interface) and the opposite interface is distinct (the unique interface) (**Fig. 5a; Supplementary Movie 4**). Across the unique interface, the relative displacement between the two assemblies is most pronounced, reaching a maximum of ∼12 Å.

**Figure 5.**
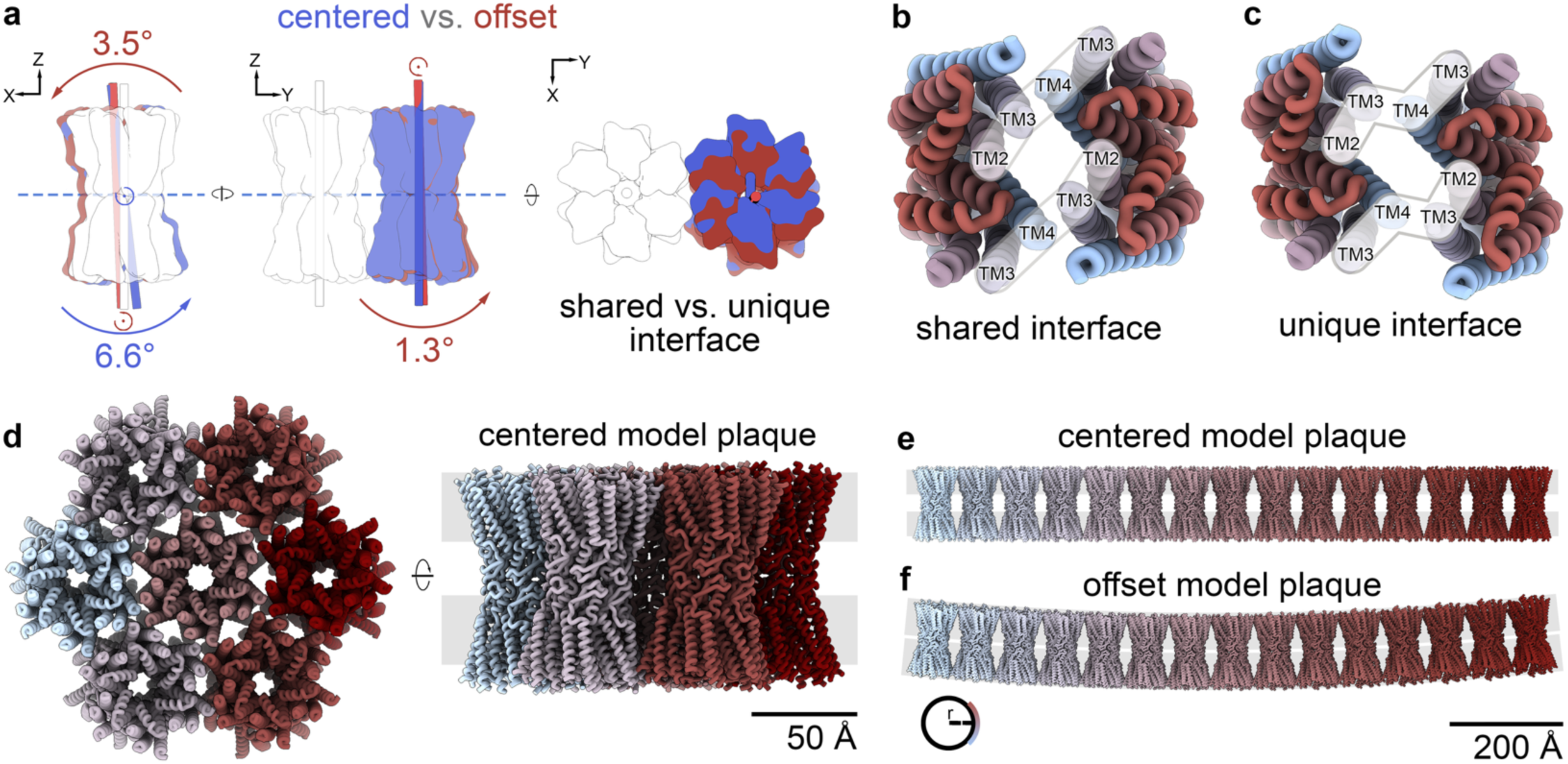
Centered and offset dual-channel geometries generate distinct packing interactions and long-range trajectories. **a,** Quantification of relative channel orientation in the centered and offset dual-channel configurations. One channel from each pair (white) was aligned to a common reference frame (pore axis along *z*), and the orientation of the partner channel (blue and red, respectively) was compared by projection into the XZ and YZ planes. The centered assembly exhibits a 6.6° rotation in the XZ plane about an axis passing through the assembly midline (blue dashed line), resulting in equivalent cytosolic-facing interfaces on both sides (two shared interfaces). The offset assembly exhibits a 3.5° rotation in XZ (centered ∼100 Å below the midline) and a 1.3° rotation in the YZ plane, producing nonequivalent cytosolic-facing interfaces (one shared and one unique interface). **b,** Top view of the shared interface highlighting transmembrane helix alignment across the interface. TM2 and TM3 from one monomer align with TM4 and TM3 from the opposing monomer to form an approximately linear arrangement. **c,** Top view of the unique interface, where the corresponding helices adopt a shifted geometry that produces a broken (zig-zag) alignment across the interface. **d,** Applying C6 symmetry to the centered dual-channel geometry generates a compact, plaque-like model containing seven channels (scale bar, 50 Å). e, Linear propagation of the centered shared interface produces an approximately straight chain of channels (scale bar, 200 Å). **f,** Linear propagation using the offset geometry produces a gradually curving chain with a radius of curvature (r) of ∼560 nm (scale bar, 200 Å). These propagated arrangements are geometric extrapolations of the observed interfaces and are not intended as direct reconstructions of native plaques.

To further quantify these orientation differences, one channel was aligned from each paired assembly to a common reference frame with the pore axis oriented along *z* (**Fig. 5a**, *white*). The partner channel in the centered and offset assemblies was then compared to this reference and relative rotations were described by their projections in the XY, XZ, and YZ planes (**Fig. 5a**, *red and blue*). In the centered assembly, the partner channel exhibits a 6.6° rotation in the XZ plane about a midline passing through the assembly, yielding equivalent displacements of ∼10 Å at the two cytosolic faces and therefore two shared interfaces. No detectable rotation is observed in the YZ plane.

In the offset assembly, the partner channel exhibits a distinct combination of rotations: a 3.5° rotation in the XZ plane about a point located ∼100 Å from the assembly midline, together with a 1.3° rotation in the YZ plane (**Fig. 5a**). This off-center rotation produces nonequivalent cytosolic-facing interfaces, generating one interface proximal to the effective rotation axis (the ‘shared’ interface) and a unique interface at the opposite face.

These geometric differences alter the local alignment of the TMs across the channel–channel interface. At the shared interface, TM2 and TM3 from one monomer align with TM4 and TM3 from the opposing monomer in an approximately linear arrangement (**Fig. 5b**). In contrast, the unique interface adopts a shifted configuration in which the corresponding helices trace a zig-zag arrangement due to the relative offset between channels (**Fig. 5c**).

Finally, to explore how the observed interface geometries could influence higher-order organization, we performed a geometric propagation analysis based on the centered and offset interfaces. Importantly, these propagated arrangements are intended as geometric extrapolations of the observed interfaces, rather than direct reconstructions of native plaques. Applying C6 symmetry to the centered assembly yields a compact, plaque-like model containing seven channels and twenty-four shared interfaces (**Fig. 5d**). Linear propagation of the shared interface produces an approximately straight chain of channels (**Fig. 5e**). In contrast, because the offset assembly contains two nonequivalent interfaces, it cannot be propagated into a C6-symmetric ring; however, stepwise linear propagation using the shared interface generates a gradually curving chain, with a radius of curvature of ∼560 nm (**Fig. 5f**). Notably, inter-channel center-to-center spacing is similar for the centered and offset assemblies (74.1 Å and 74.5 Å, respectively), indicating that the primary difference between these packing modes is geometric orientation rather than overall separation.

## DISCUSSION

Understanding how gap junction channels assemble into plaques and how this high-order organization shapes channel function is essential for understanding intercellular signaling in tissues and how its dysregulation contributes to disease. In the avascular eye lens, Cx46/50 gap junctional plaques support the lens microcirculatory system by enabling long-range intercellular flow of nutrients and antioxidants and maintaining ionic and metabolic homeostasis^48^. Plaque organization in lens fiber cells is tightly coupled to lens physiology and deteriorates with age alongside reduced gap junctional coupling and cataract-associated dysfunction^49,50^.

To gain mechanistic insight into this problem we establish a reconstitution strategy that captures multiple native lens Cx46/50 intercellular channels within a shared lipid membrane scaffold, enabling high-resolution structural interrogation of channel–channel organization in a defined bilayer environment. By resolving dual-channel assemblies in two distinct geometries, we identify a central principle of multi-channel assembly – neighboring channels associate without ordered protein–protein contacts across structured regions. Instead, the inter-channel space is occupied by ordered lipid features, indicating that channel packing in these assemblies is lipid mediated. This principle offers a direct explanation for how plaques can achieve dense lateral packing while retaining local variability in relative channel orientation/spacing, a hallmark of plaques observed by electron microscopy and two-dimensional crystallography^37,39,51^. In addition, the large dataset acquired for this heterogeneous ensemble of channel assemblies enabled refinement of the single-channel Cx46/50 reconstruction to 1.8 Å, providing an exceptional baseline for defining the conductive open-state and internal reference for assessing lipid features that emerge specifically in dual-channel assemblies.

### Gap junctional inter-channel assembly mediated by leaflet-specific lipid ordering

A second conclusion from these structures is that plaque-like packing can stabilize discrete lipid populations at the channel–channel interface, including features not resolved in single-channel reconstructions. In the centered dual-channel assembly, ordered extracellular leaflet lipids extend across the entire inter-channel gap and adopt a continuous hexagonal packing arrangement – including headgroup features and putative protein–headgroup interactions not readily resolved in single-channel reconstructions. Strikingly, we also resolve lipid headgroup features in the intracellular-facing leaflet at the shared interface, another feature not observed in our single-channel reconstructions of Cx46/50 under comparable non-gating conditions^16^. Together, these observations indicate that channel proximity can increase bilayer order locally and reveal leaflet-specific differences in lipid stabilization, with extracellular leaflet acyl chains and interfacial headgroups exhibiting distinct patterns of resolvability. More broadly, the presence of ordered lipids spanning the full interface suggests that local packing constraints may be influenced by the composition of the bilayer itself, potentially encoding allowable channel spacing through the number and arrangement of specific stabilized lipids.

### Multiple interface geometries support plasticity of higher-order assembly

The centered and offset dual-channel assemblies define two interface geometries with distinct symmetry properties at the cytosolic faces. The centered assembly yields equivalent shared interfaces on both sides of the paired channels, whereas the offset assembly contains one shared interface and one unique interface. These two arrangements likely represent only a subset of an accessible landscape of possible relative orientations that can occur when channel–channel packing is coupled through a deformable membrane bilayer. In this context, conformational heterogeneity is an intrinsic feature of lipid-mediated assembly. Geometric propagation of the observed interfaces illustrates potential consequences of these packing modes, with the centered interface supporting straighter trajectories and the offset arrangement producing gradual curvature. Indeed, early models of curved gap junction plaques show similar architecture to the offset configuration^52^. While these propagated models are intended as geometric extrapolations rather than true reconstructions of native plaques, they provide an intuitive link between local interface geometry and mesoscale organization, including the possibility that mixtures of related interfaces could contribute to heterogeneous plaque morphology *in vivo*.

### Connecting reconstituted membrane assemblies to in situ plaque architecture

A key question is how the channel spacing and lipid organization captured in this simplified reconstitution system relate to native plaques. In our dual-channel assemblies, channels pack at ∼74 Å center-to-center, closely matching reconstituted 2D protein–lipid crystals^7,40,53^ but tighter than many measurements from isolated native plaques and *in situ* studies (∼80–90 Å)^34,35,39,51,54,55^. This disparity suggests that native membrane composition, associated scaffolding proteins, and/or membrane ultrastructure tune the energetics of channel packing, thereby shifting the preferred lattice spacing and degree of order. This is likely particularly relevant in lens fiber cells, where plaques are often quasi-ordered^55^ and the membrane is highly specialized, enriched in cholesterol and sphingolipids, and remodeled with age^49,56–58^. Notably, the absence of ordered protein–protein contacts in our structures despite relatively tight packing supports the idea that close plaque-like organization can be achieved primarily through lipid-mediated coupling rather than extensive interfaces between structured channel domains.

Recent advances in cryo-electron tomography (cryo-ET) are beginning to bridge the long-standing gap between atomic-resolution connexin structures and mesoscale plaque organization in native membranes^54,59^. For example, recent *in situ* reconstructions of connexin plaques containing Cx43 highlight the potential of cryo-ET to define plaque architecture directly within cells^54^. At present, however, subtomogram averages of gap junctions have not reached the resolution needed to resolve specific lipid interactions or to unambiguously visualize critical domain-specific features, such as the NT gate, that would report on the channel functional state. Even so, these *in situ* structural characterizations provide an essential framework for understanding how plaques assemble in their physiological context and motivate complementary high-resolution approaches. Together, *in situ* cryo-ET and reductionist single-particle reconstructions offer a convergent path toward a molecular description of plaque formation that spans scales from membrane-encoded interfaces to native ultrastructure.

### Interstitial lipid occupancy as a potential link between packing and function in Cx46/50

Although both dual-channel assemblies adopt the stabilized open state, the centered shared interface uniquely resolves an intercalated interstitial lipid (IL) whose headgroup engages the open-state NT, a feature not detected in our single-channel reconstructions in the absence of gating stimuli. This observation suggests that channel–channel packing can bias lipid occupancy near the NT region without necessarily shifting the channel into a closed conformation. Notably, recent pH-gating structures of Cx46/50 reveal lipid infiltration into the pore concomitant with NT displacement and stabilization of closed conformations under acidification^17^, providing an experimentally observed endpoint for a lipid-engaged gating pathway.

Given its position adjacent to the NT gating domain and its open-state dependent interactions with Q15 (N15 in Cx50) and E12/E16, this IL site offers a plausible structural route by which higher-order organization could modulate pore-proximal lipid occupancy and thereby tune gating energetics, potentially destabilizing the open-state and/or facilitating access to lipid-mediated gating transitions. More broadly, these observations are consistent with the idea that plaque-like packing could potentiate lipid engagement along the pore under appropriate stimuli or membrane conditions, offering potential insight into why junctional conductance in tissues is often far lower than predicted from plaque size and unitary conductance. Future studies that vary lipid composition and apply controlled gating stimuli of plaque-like assemblies should help define how membrane environment and higher-order organization jointly tune lipid engagement near the pore.

### Limitations and outlook

This work uses a simplified MSP/DMPC system to capture multi-channel organization at high resolution, enabling direct visualization of lipid-mediated interfaces and pore-proximal lipid occupancy. At the same time, native gap junction plaques incorporate complex lipid mixtures, additional membrane proteins, and cytoskeletal and scaffolding elements that are not represented here. In the lens, where plaques form in a uniquely cholesterol-rich and age-remodeled membrane environment, extending these approaches to complex lipid mixtures and native-like membrane systems will be particularly important for understanding how plaque organization evolves with aging and cataractogenesis. In addition, native lens Cx46/50 purified from bovine lens core tissue exhibit post-translational processing, including C-terminal truncation^60^, which may influence plaque organization *in vivo* in ways that may be difficult to capture by single-particle approaches^61^. These limitations motivate future extensions in which multi-channel assemblies are reconstituted into more native-like lipid mixtures, combined with controlled modulation of pH or calcium, and integrated with cryo-ET studies of native membranes. Together, such approaches should help establish how lipid-mediated coupling, disordered cytosolic regions, and cellular ultrastructure collectively shape plaque architecture and function across tissues.

## ACKNOWLEDGEMENTS

We thank Dr. Claudia López and the staff of the OHSU Multiscale Microscopy Core for training and instrumentation access. We also acknowledge computational support from the OHSU Advanced Computing Center (NIH S10OD034224) and cryo-EM instrumentation and user support from the Pacific Northwest Cryo-EM Center (R24GM154185). This work was supported by NIH R35GM124779 (S.L.R.) and NIH T32EY023211 (C.S.G.).

## AUTHOR CONTRIBUTIONS

S.A.S. performed sample preparation and cryo-EM grid preparation. J.M.J. performed cryo-EM grid preparation and screening. J.M.J. and J.B.M. collected cryo-EM data. J.M.J. generated the preliminary dual-channel reconstruction. J.B.M. performed image processing and atomic modeling for the single-channel reconstruction. C.S.G. performed image processing and atomic modeling for the dual-channel reconstructions, conducted comparative structural analyses, and drafted the manuscript. S.L.R. supervised the project and revised the manuscript. All authors contributed to the final version of the manuscript.

## MATERIALS AND METHODS

### MSP expression and purification

A His-tagged MSP1E1 expression plasmid was obtained from Addgene^62^. MSP1E1 was expressed in *E. coli* as previous described^16,17^, and cell pellets were suspended in lysis buffer (40 mM Tris pH 7.4, 1% Triton X-100, 1 mM PMSF, 150 mM NaCl) and stored at −80° C until use. MSP1E1 was purified as previously described^16,42^, with minor modifications. Briefly, frozen cell pellet suspension (4 g wet cells) was thawed on ice, supplemented with 100 µM PMSF, and lysed by sonication. Lysate was clarified by ultracentrifugation (110,000×g, 30 min, 4° C), and the supernatant was applied to 5 mL Ni-NTA resin pre-equilibrated in binding buffer (40 mM Tris [pH 7.4], 300 mM NaCl) using a gravity column. Binding was performed in batch for 1 h at 4° C with gentle rocking. The resin was then washed sequentially with 5 column volumes (CV) of binding buffer, followed by 5 CV of binding buffer supplemented with 1% Triton X-100 and 1 mM PMSF, 5 CV of binding buffer supplemented with 50 mM sodium cholate, and 5 CV of binding buffer supplemented with 50 mM imidazole. Finally, MSP1E1 was eluted with 5 CV of binding buffer supplemented with 500 mM imidazole.

Eluted protein was concentrated to 5.8 mg mL^-1^ using a 5 kDa MWCO centrifugal concentrator (Vivaspin 6, Sartorius) and filtered (0.22 µm, Millipore). The sample was further purified by size-exclusion chromatography (SEC) on an EnRich 70 (Bio-Rad) column equilibrated in running buffer (20 mM HEPES [pH 7.4], 150 mM NaCl) using an FPLC system (Bio-Rad NGC) at 4° C. Peak fractions were pooled and concentrated to 300 µM using a 5 kDa MWCO concentrator (Vivaspin 6). Aliquots were flash-frozen in liquid nitrogen and stored at −80° C.

### Cx46/50 purification and reconstitution into lipid nanodiscs

Fresh bovine eyes were obtained from Nebraska Scientific (Omaha, NE). Lenses were surgically isolated and stored at −80 °C until use. Stripped membrane was prepared from lens core tissue, which is enriched in C-terminally truncated Cx46/50 gap junctions, as previously described^15,16^. Stripped membranes were stored at −80° C in storage buffer (10 mM Tris [pH 8.0], 2 mM EDTA, 2 mM EGTA) at 2 mg mL^-1^ total protein (BCA assay, Pierce).

For solubilization, stripped membranes were thawed and mixed 1:1 (v/v) with solubilization buffer (10 mM Tris [pH 8.0], 2 mM EDTA, 2 mM EGTA, 2% n-decyl-β-D-maltoside (DM; Anatrace)). The mixture was incubated at 37° C for 30 min on a rotator. Insoluble material was removed by ultracentrifugation (110,000×g, 30 min, 4° C). The supernatant was filtered (0.22 µm, Millipore) and applied to an anion-exchange column (UnoQ, Bio-Rad). Protein was eluted using a 22 CV gradient from Buffer A (10 mM Tris [pH 8.0], 2 mM EDTA, 2 mM EGTA; 0.02% DM) to Buffer B (Buffer A supplemented with 350 mM NaCl). Peak fractions were pooled and concentrated using a 50 kDa MWCO centrifugal concentrator (Vivaspin 6) to a final protein concentration of 150 µM.

Cx46/50 was reconstituted into nanodiscs as previously described^16^ with minor modifications detailed here. DMPC stocks in chloroform were dried under nitrogen and placed under vacuum overnight. Dried lipid film was resuspended to 25 mM in lipid solubilization buffer (20 mM HEPES [pH 7.4], 150 mM NaCl, 2 mM EDTA, 2 mM EGTA, 5% DM) and solubilized by bath sonication at 37° C. Purified Cx46/50 was incubated with solubilized DMPC at a 0.6:90 molar ratio (protein:lipid) with gentle agitation for 1 h at 25° C. MSP1E1 was then added to achieve a final molar ratio of 0.6:1:90 (Cx46/50:MSP1E1:DMPC), followed by an additional 15 min incubation at 25° C with gentle agitation.

Detergent was removed by addition of SM-2 Bio-Beads (Bio-Rad) at a 30:1 bead:detergent mass ratio and incubation with rotation overnight at 25° C. Fresh beads were replaced (30:1 bead:detergent, w/w) for an additional 1 h at 25° C. Following bead removal, insoluble material was pelleted (16,000×g, 30 min, 4° C), and the supernatant was filtered (0.22 µm) and subjected to SEC on a Superose 6 Increase 10/300 GL column (Cytiva) equilibrated in running buffer (20 mM HEPES [pH 7.4], 150 mM NaCl, 2 mM EDTA, 2 mM EGTA) to separate Cx46/50-containing nanodiscs from empty nanodiscs. Fractions corresponding to 10.5–13.5 mL elution volume were pooled and concentrated at 4° C using a 50 kDa MWCO concentrator (Vivaspin 6) at 500×g to 9.7 µM particle concentration (estimated λ_280_ ∼ 631,320 M^-1^cm^-1^).

### Negative stain electron microscopy

Nanodisc-solubilized Cx46/50 was prepared for negative-stain electron microscopy as previously described^15,16^. Briefly, 3 µL of sample was applied to a glow-discharged continuous carbon copper grid (Ted Pella, G400). After blotting with filter paper, grids were washed three times in SEC running buffer, blotting after each wash. Grids were then stained twice with freshly prepared uranyl formate (0.75% w/v; SPI-Chem), blotted, and dried under laminar airflow. Data were collected on a 120 kV TEM (Tecnai T12, FEI) equipped with a 2k CCD camera (Eagle, FEI) at a nominal magnification of 49,000× (4.401 Å px^-1^), using ∼1.5 µm defocus.

### Cryo-EM specimen preparation and data collection

Freshly prepared Cx46/50 nanodisc sample was mixed in a PCR tube with size-exclusion running buffer and DMSO to a final concentration of 1% (v/v), then incubated for 1 h at 25° C. For vitrification, 3 µL of sample (∼3–4 µM particle concentration) was applied to glow-discharged Quantifoil R2/1 Cu 300 mesh holey carbon grids and plunge-frozen in liquid ethane using a Vitrobot Mark IV (Thermo Fisher Scientific) operated at 100% humidity and 23° C with a 5 s blot time. Grids were stored in liquid nitrogen prior to imaging.

Cryo-EM data were initially screened at the OHSU Multiscale Microscopy Core (MMC) on a 200 kV Glacios (Thermo Fisher Scientific), with final high-resolution data collected at the Pacific Northwest Cryo-EM Center (PNCC) on a 300kV Titan Krios G4 (Thermo Fisher Scientific) operated using SerialEM. A total of 7,884 movies were recorded on a Gatan K3 direct electron detector at 105,000× nominal magnification in super-resolution mode (0.83 Å physical pixel size; 0.41 Å super-resolution pixel size), with an energy filter (BioContinuum, Gatan) slit width of 10–20 eV. Movies were recorded as 46-frame exposures with a total dose of 40 e⁻ Å^-2^ and a nominal defocus range of −1.8 to −0.5 µm (**Supplementary Table 1**).

### Image processing of the single-channel assembly

All image processing was performed in CryoSPARC (v4.7.1)^63^. Movies were motion-corrected using Patch Motion Correction with Fourier cropping to the physical pixel size (2× binning). CTF parameters were estimated using Patch CTF Estimation. Micrographs were denoised using the Micrograph Denoiser job after training on 100 representative micrographs. Particles were initially picked with a blob picker on denoised micrographs, yielding 7,652,025 particles. Particles were extracted with a box size of 432 pixels and Fourier-cropped to 84 pixels (4.27 Å pixel ^-1^) to expedite initial 2D classification. The initial particle set was culled to 807,5151 particles after two rounds of classification by keeping only single channel classes. A subset of 315,338 particles representing the highest detail 2D classes was used for ab initio reconstruction to generate initial 3D references.

For 3D refinement, the 807,515-particle stack was re-extracted with a box size of 384 pixels and Fourier-cropped to 192 pixels. An initial non-uniform refinement without imposed symmetry produced a 3D reconstruction at 3.4 Å nominal resolution. To further remove lower-quality particles, asymmetric heterogeneous refinement (five classes) was performed, yielding one high-quality class containing 430,947 particles. Non-uniform refinement of this class with D6 symmetry imposed yielded a 3.4 Å reconstruction (Nyquist limit at this binning). Particles were then re-extracted at a box size of 384 pixels without additional Fourier cropping, and non-uniform refinement with D6 symmetry imposed reached 2.04 Å nominal resolution. Reference-based motion correction improved the reconstruction to 1.94 Å, followed by global and local CTF refinement, yielding a final reconstruction at 1.80 Å from 402,098 particles with D6 symmetry imposed.

A final round of heterogeneous refinement (four classes; D6 symmetry) produced two high-quality, high-population classes and two lower-quality classes. The combined high-quality classes (approximately 355,000 particles) refined to 1.79 Å with D6 symmetry imposed. Further classification did not improve the reconstruction and generally reduced the achieved resolution, consistent with resolution-limited behavior for the available particle count (ResLog analysis).

### Dual-channel cryo-EM image processing

Particles were picked by template matching using a preliminary dual-channel reconstruction generated from a separate dataset^17^ (EMPIAR-13131, derived by 2D classification followed by non-uniform refinement) as the template in the Template Picker job. In total, 1,977,900 particles were extracted at 4.3 Å pixel^-1^ and subjected to a single round of 2D classification (200 classes) to remove non-particle and low-quality classes, yielding 1,345,806 particles.

### Initial 3D classification

Particles retained after 2D classification were subjected to *ab initio* reconstruction with four classes. One class corresponding to a single-channel reconstruction and one class resembling a dual-channel reconstruction were retained, while classes lacking interpretable features were discarded. The retained particles were subjected to a second *ab initio* reconstruction (four classes), resulting in one class resembling a dual-channel reconstruction (retained) and three classes discarded. Particles corresponding to the single-channel class from the first *ab initio* job were refined independently using homogeneous refinement with D6 symmetry to generate a single-channel reference volume for downstream use.

Following this *ab initio* 3D classification workflow, the dual-channel particle stack still exhibited substantial contamination by single-channel particles. To remove these contaminants, the particle stack was subjected to one round of “baited” heterogeneous refinement, in which particles were classified against two competing reference volumes. The references used were the dual-channel template employed for template-based picking (EMPIAR-13131) and the D6-symmetrized single-channel reconstruction generated above. Particles assigned to the single-channel class were discarded, and the remaining 262,017 particles assigned to the dual-channel class were retained for further processing.

### Resolving dual-channel configurational heterogeneity by 3D variability analysis

Despite extensive removal of single-channel contaminants, the dual-channel reconstruction following ‘baited’ heterogeneous refinement showed asymmetric resolvability of the two channels, suggesting additional heterogeneity. To address this, particles and the consensus volume were subjected to two rounds of 3D variability analysis (3DVA, 6 Å filter resolution). The dominant component (PC 1) separated single- and dual-channel particles. The dual-channel half of the component (133,365 particles) was pooled, re-extracted (280 pixel box, 1.28 Å pixel^-1^), and refined by non-uniform refinement to 2.95 Å but still exhibited asymmetric channel resolvability.

A second round of 3DVA identified a principal component (PC 1) for which both endpoints yielded reconstructions with two well-resolved channels. Particles from each end of the component were pooled and refined independently by non-uniform refinement, producing two C1 reconstructions corresponding to the centered and offset dual-channel configurations (centered: 3.37 Å, 64,687 particles; offset: 4.28 Å, 91,636 particles).

Both configurations were then subjected to additional rounds of heterogeneous refinement and non-uniform refinement to remove low-quality particles, followed by global and local CTF refinement, local motion correction, and local resolution filtering. The final centered reconstruction was refined to 2.88 Å with D2 symmetry (64,687 particles). The final offset reconstruction was refined to 3.43 Å with C2 symmetry (53,547 particles). For each configuration, corresponding C1 reconstructions were also generated with local filtering for comparison.

### Atomic modeling and validation

Previously published ovine Cx46 (PDB 7JKC) and Cx50 (PDB 7JJP) models were used as starting coordinates^16^ and sequence-mutated to bovine in UCSF ChimeraX^64^. For the single-channel assembly, model building was performed in Coot^65^ using the D6-symmetrized map, treating a single connexin subunit as the asymmetric unit. The full dodecameric channel was generated using non-crystallographic symmetry (NCS) operators in Phenix^66^. For the paired-channel reconstructions, models were built in Coot into the D2-symmetrized centered map (hemichannel asymmetric unit) and the C2-symmetrized offset map (dodecameric channel asymmetric unit), and full paired-channel assemblies were generated using NCS operators in Phenix.

Iterative cycles of manual rebuilding in Coot and real-space refinement in Phenix were performed until convergence, with model quality assessed using MolProbity^67^. Because Cx46 and Cx50 are not distinguishable in the cryo-EM density maps, Cx46 was refined first as the archetype model and then converted to Cx50 by sequence mutation and refined using the same procedures. DMPC lipids were modeled where supported by density, with acyl chains trimmed to match map features. Water molecules were placed only when supported by density in both half maps and consistent with local hydrogen-bonding chemistry. Structural visualization and analysis were performed in UCSF ChimeraX.

### Dual-channel orientation analysis

Relative orientations of dual-channel models were quantified using UCSF ChimeraX (v1.9)^64^. For each intercellular channel, the pore axis was defined by placing centroids at the centers of mass of the two hemichannels and connecting these centroids with a vector. Relative channel orientations were computed by aligning one channel from each paired assembly to a common reference frame and calculating angles between the resulting pore-axis vectors after projection onto the XY, XZ, and YZ planes. Angle calculations were performed in Python 3 (repository: https://github.com/reichow-lab/Multichannel-Analysis-Scripts).

### AI-assisted technologies

During the development of this work, ChatGPT (OpenAI) was used for help in writing code and to revise portions of the text to improve clarity. After using this tool for coding, synthetic datasets were generated and used for testing and validation. The authors reviewed and edited any content that was generated with this tool and take full responsibility of the content of this publication.

## DATA AVAILABILITY

Cryo-EM maps have been deposited in the Electron Microscopy Data Bank (EMDB) under accession codes EMDB-XXXXX (single-channel), EMDB-XXXXX (centered dual-channel), and EMDB-XXXXX (offset dual-channel). Atomic coordinates have been deposited in the Protein Data Bank (PDB) under accession codes XXXX (Cx46, single-channel), XXXX (Cx46, single-channel), XXXX (Cx46, centered dual-channel), XXXX (Cx46, offset dual-channel), XXXX (Cx50, centered dual-channel), and XXXX (Cx50, offset dual-channel). The original multi-frame micrographs have been deposited in the Electron Microscopy Public Image Archive (EMPIAR) under accession code EMPIAR-XXXXXX.

## CODE AVAILABILITY

Code for the analysis of relative channel orientations is available on the Reichow Lab GitHub repository (https://github.com/reichow-lab/Multichannel-Analysis-Scripts).

## SUPPLEMENTARY INFORMATION

## SUPPLEMENTARY TABLES AND FIGURES

**Supplementary Table 1:** Cryo-EM data collection, refinement, and validation statistics.

| Conformation | Centered |  | Offset |  | Single Channel |  |
| --- | --- | --- | --- | --- | --- | --- |
| Data Collection and Processing |  |  |  |  |  |  |
| Magnification | 105,000x |  | 105,000x |  | 105,000x |  |
| Voltage (kV) | 300 |  | 300 |  | 300 |  |
| Electron Exposure (e<sup>-</sup> / Å<sup>2</sup>) | 40 |  | 40 |  | 40 |  |
| Defocus Range (μm) | -0.8 to -0.5 |  | -0.8 to -0.7 |  | -0.8 to -0.9 |  |
| Pixel Size (Å) | 0.41 |  | 0.41 |  | 0.41 |  |
| Symmetry Imposed | D2 |  | C2 |  | D6 |  |
| Initial particle Images | 1,354,805 |  | 1,354,805 |  | 806,504 |  |
| Final Particle Images | 64,687 |  | 53,547 |  | 355,000 |  |
| Map Resolution | 2.88 |  | 3.43 |  | 1.79 |  |
| FSC Threshold | 0.143 |  | 0.143 |  | 0.143 |  |
| Map Resolution Range (Å) | 1.78 to 30 |  | 1.76 to 30 |  | 1.56 to 22 |  |
| Model Refinement |  |  |  |  |  |  |
| Isoform | Cx46 | Cx50 | Cx46 | Cx50 | Cx46 | Cx50 |
| Initial Model (PDB ID) | 7JKC | 7JJP | 7JKC | 7JJP | 7JKC | 7JJP |
| Model Resolution (Å) | 2.9 | 2.9 | 3.5 | 3.5 | 1.8 | 1.8 |
| FSC Threshold | 0.5 | 0.5 | 0.5 | 0.5 | 0.5 | 0.5 |
| Model resolution range (Å) | 2.4 to 2.9 | 2.4 to 2.9 | 2.8 to 3.5 | 2.8 to 2.5 | 1.6 to 1.8 | 1.6 to 1.8 |
| Map sharpening B-factor (Å<sup>2</sup>) | 66 | 66 | 57.7 | 57.7 | 53.3 | 53.3 |
| Model Composition |  |  |  |  |  |  |
| Non-hydrogen atoms | 39,136 | 38,872 | 37,176 | 36,932 | 44,304 | 44,088 |
| Protein Residues | 4,416 | 4,416 | 4,416 | 4,416 | 2,292 | 2,292 |
| Ligands | 312 | 312 | 116 | 116 | 648 | 648 |
| B factors (Å<sup>2</sup>) |  |  |  |  |  |  |
| Protein | 53.61 | 53.62 | 53.61 | 75.06 | 9.89 | 10.76 |
| Ligands | 57.37 | 57.33 | 37.6 | 39.95 | 19.81 | 17.15 |
| RMS deviations |  |  |  |  |  |  |
| Bond lengths | 0.01 | 0.011 | 0.008 | 0.005 | 0.004 | 0.006 |
| Bond angles | 1.293 | 1.373 | 1.2 | 1.071 | 0.784 | 0.812 |
| Validation |  |  |  |  |  |  |
| MolProbity Score | 1.48 | 1.49 | 1.6 | 1.54 | 1.14 | 1.19 |
| Clash score | 4.52 | 4.76 | 5.73 | 6.82 | 2.65 | 3.55 |
| Poor rotamers (%) | 0.41 | 1.2 | 0.92 | 0.62 | 1.18 | 1.14 |
| Ramachandran Plot |  |  |  |  |  |  |
| Favored (%) | 95.93 | 96.11 | 95.83 | 97.08 | 97.86 | 98.4 |
| Allowed (%) | 4.07 | 3.89 | 4.17 | 2.92 | 2.14 | 1.6 |
| Disallowed (%) | 0 | 0 | 0 | 0 | 0 | 0 |

**Supplementary Figure 1:**
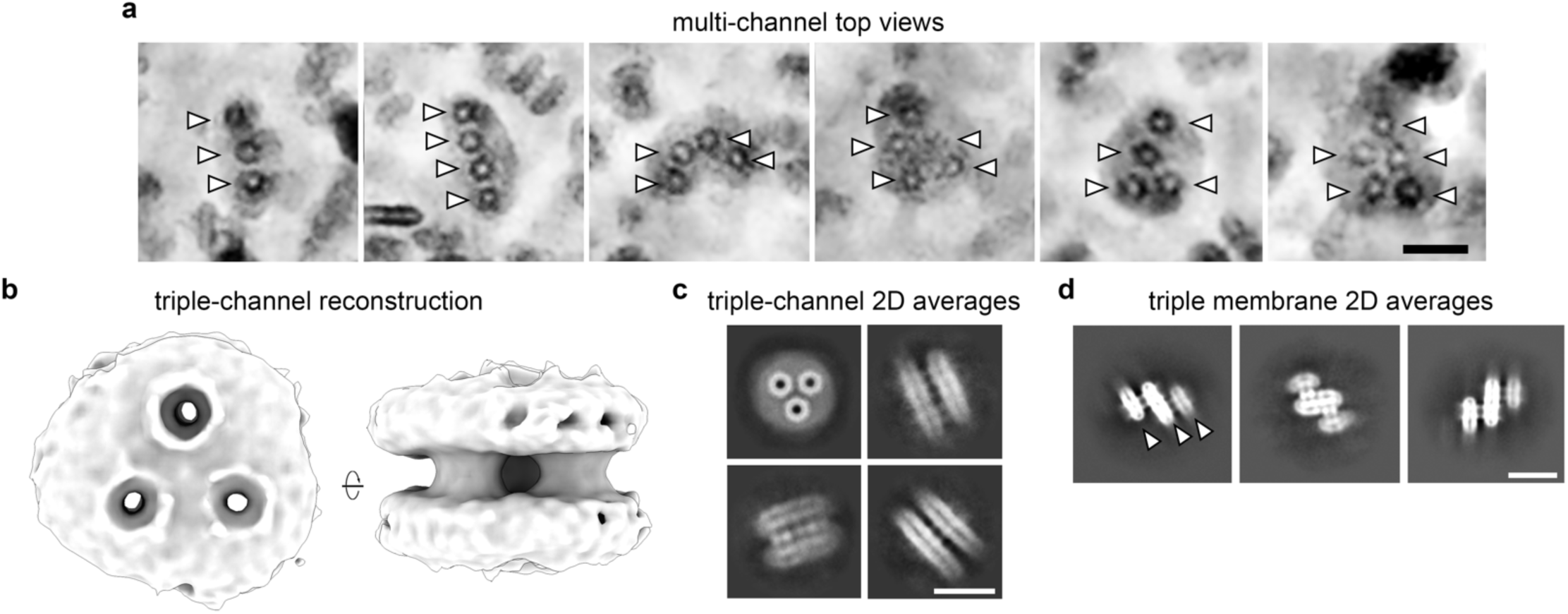
Representative high-order assemblies in the multi-channel Cx46/50 nanodisc sample. **a,** Representative denoised cryo-EM micrograph views showing membrane particles containing multiple (three or more) gap junction channels (arrowheads). Scale bar, 100 Å. **b,** *Ab initio* 3D reconstruction of a triple-channel assembly refined without imposed symmetry (C1), at ∼11.5 Å resolution. **c**, Representative 2D class averages of triple-channel assemblies. Scale bar, 100 Å. **d,** Representative 2D class averages of “triple-membrane” assemblies containing three bilayer layers (arrowheads). Scale bar, 100 Å.

**Supplementary Figure 2:**
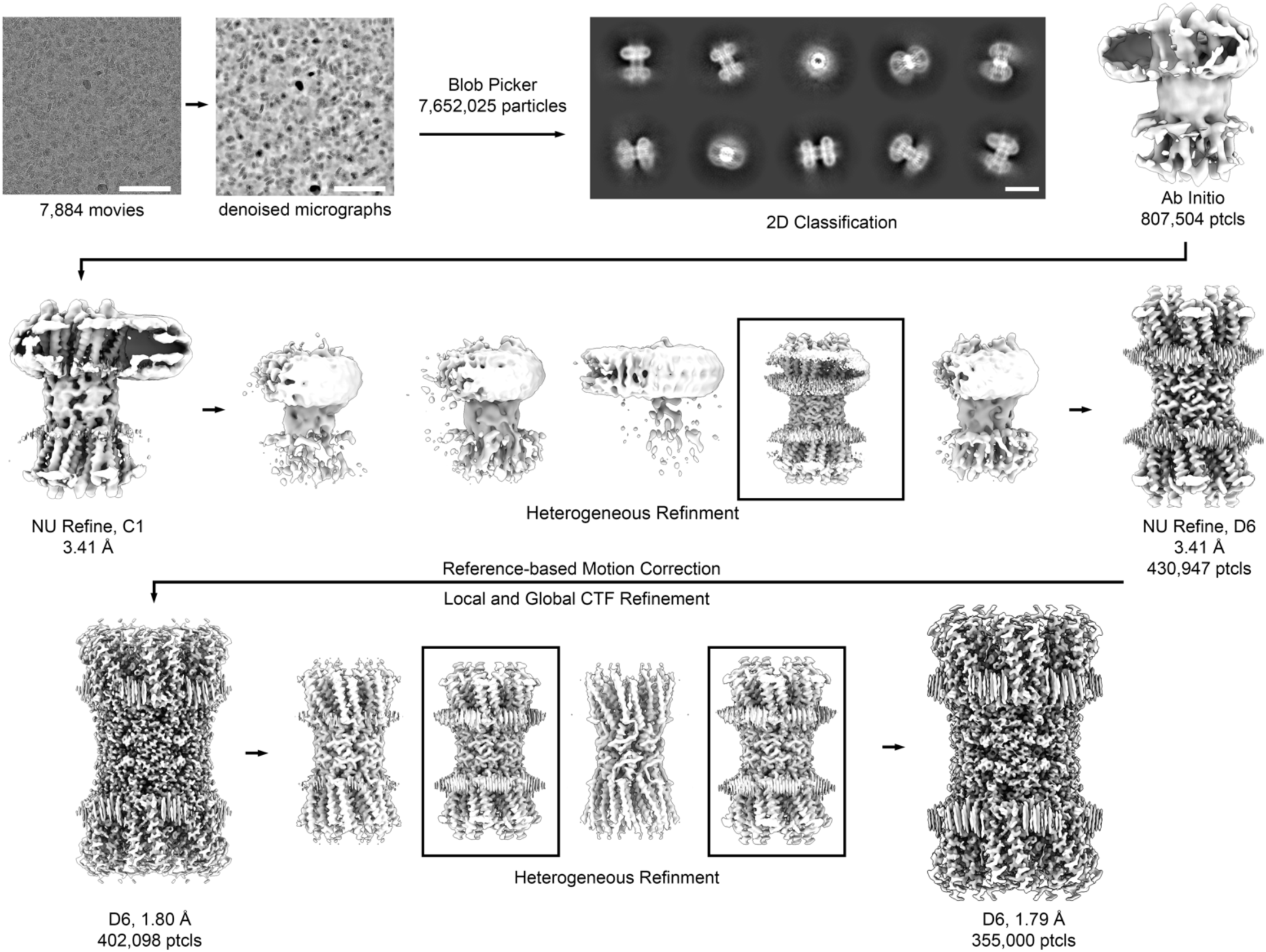
Single-channel cryo-EM processing workflow. Workflow for single-channel Cx46/50 reconstruction in CryoSPARC. Movies (7,884) were preprocessed by motion correction, CTF estimation and denoising to facilitate particle picking. Blob Picker was used for initial particle selection (7,652,025 particles), followed by 2D classification, and *ab initio* reconstruction to generate initial 3D references. Subsequent non-uniform refinement, heterogeneous refinement, reference-based motion correction, and global/local CTF refinement yielded the final D6-symmetrized reconstruction at 1.80 Å (402,098 particles) and a refined subset at 1.79 Å (355,000 particles). Scale bars: micrographs, 100 nm; 2D class averages, 100 Å.

**Supplementary Figure 3:**
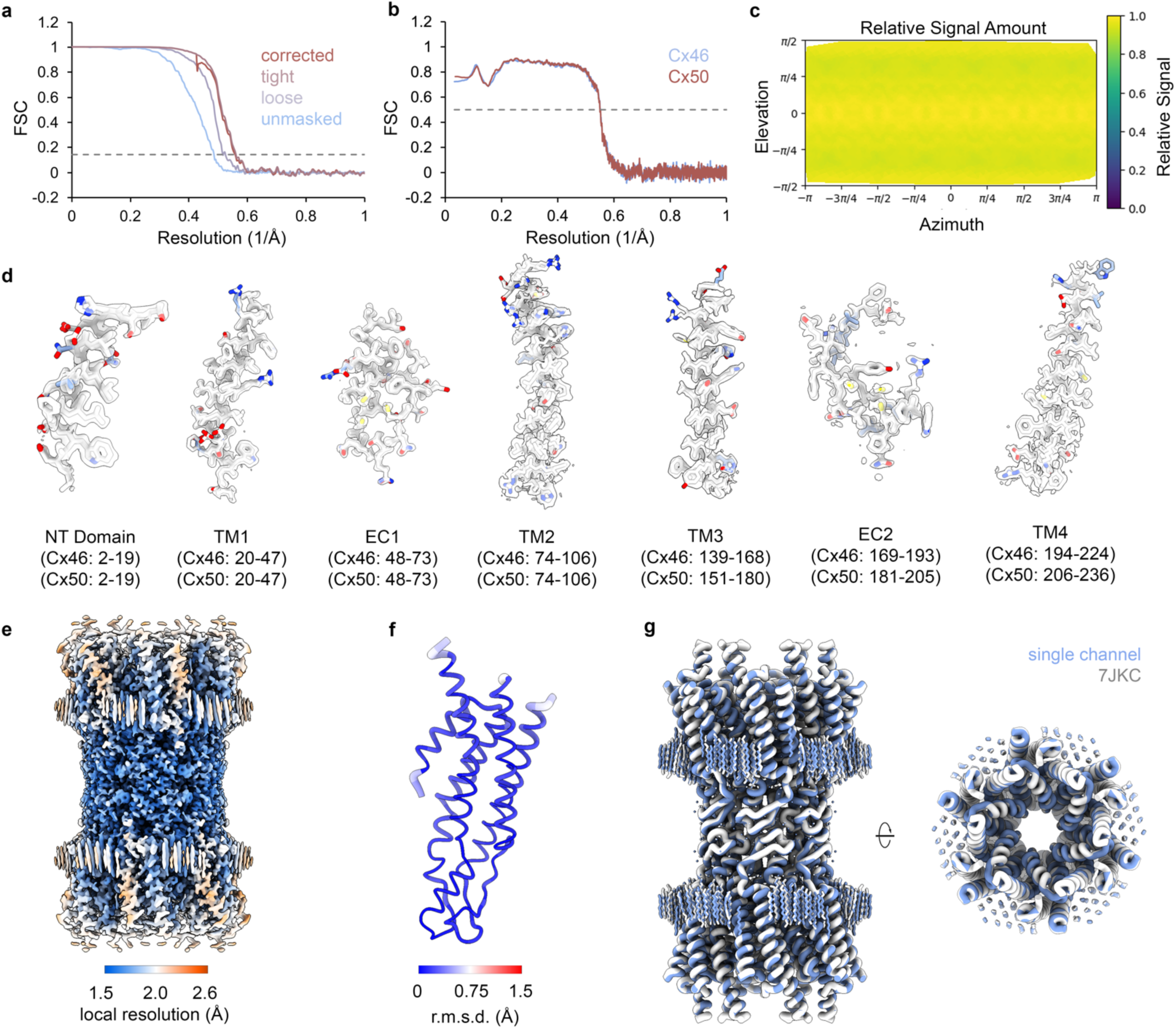
Map quality, validation, and local resolution of the single-channel reconstruction. **a,** Gold-standard Fourier shell correlation (FSC) curves for the single-channel reconstruction using different masking strategies (unmasked, loose, tight, and corrected). The dashed line indicates the FSC = 0.143 cutoff. **b,** Unmasked map-to-model FSC curves for Cx46 (blue) and Cx50 (red). The dashed line indicates the FSC = 0.5 cutoff. **c,** Azimuth–elevation plot of relative signal as a function of particle viewing direction, indicating no strong preferred orientation. **d,** Representative map-to-model fits for structured regions of the single-channel reconstruction, including the NT domain, transmembrane helices (TM1–TM4), and extracellular loops (EC1–EC2) (residue ranges indicated). Models for Cx46 and Cx50 are overlayed and colored by atom type. Cx46 – white, Cx50 – blue. **e,** Cryo-EM map colored by local resolution. Resolution is highest at the central docking domains and decreases outward radially toward the cytoplasmic regions. f, Structural alignment of a single-channel bovine Cx46 subunit from this study with the previously reported ovine Cx46 subunit (PDB 7JKC). Cα r.m.s.d. is mapped onto the model by both worm radius and color (color scale shown). **g,** Alignment of the single-channel model from this study (blue) with PDB 7JKC (white), illustrating highly similar overall architecture, including stabilized annular lipid features.

**Supplementary Figure 4:**
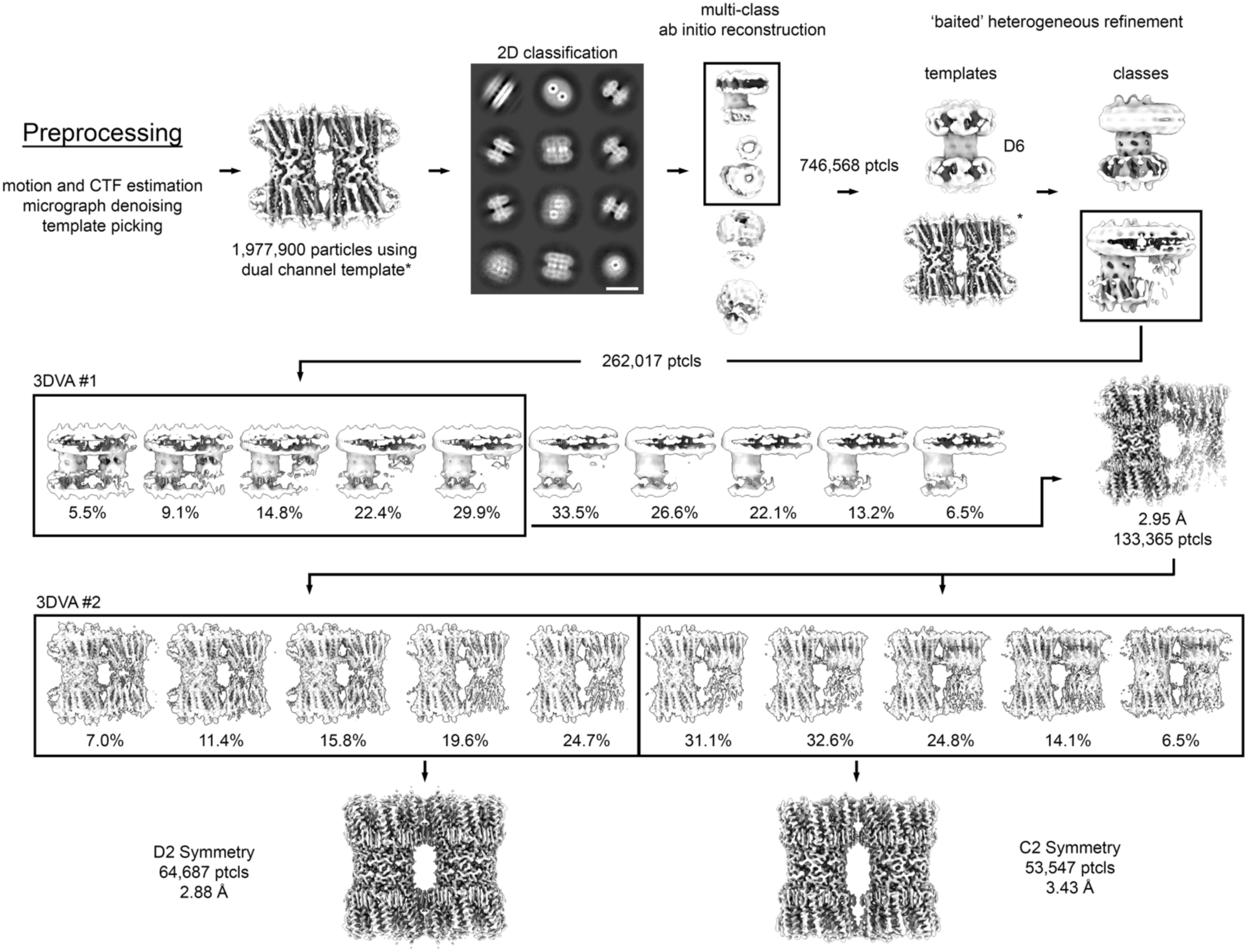
Image processing workflow for isolating centered and offset dual-channel particle sets. Overview of dual-channel image processing. Preprocessing steps (motion correction, CTF estimation, micrograph denoising) were performed as described in Supplemental Figure 2. Template picking was carried out using a preliminary dual-channel reconstruction from a prior dataset^17^ (asterisk). Following 2D classification and multi-class *ab inito* reconstruction, particles were subjected to ‘baited’ heterogeneous refinement against competing single-channel (D6) and dual-channel (C1) templates. The resulting particle stack was subjected to two rounds of 3DVA as particle curation steps. In 3DVA #1, the dominant principal component captured compositional heterogeneity and enabled removal of residual single-channel contaminants, yielding a dual-channel particle subset (133,365 particles; 2.95 Å C1 refinement). In 3DVA #2 (PC1), variability captured configurational heterogeneity and separated particles into two dual-channel configurations that were refined independently to the final reconstructions: centered (D2 symmetry, 2.88 Å; 64,687 particles) and offset (C2 symmetry, 3.43 Å; 53,547 particles).

**Supplementary Figure 5:**
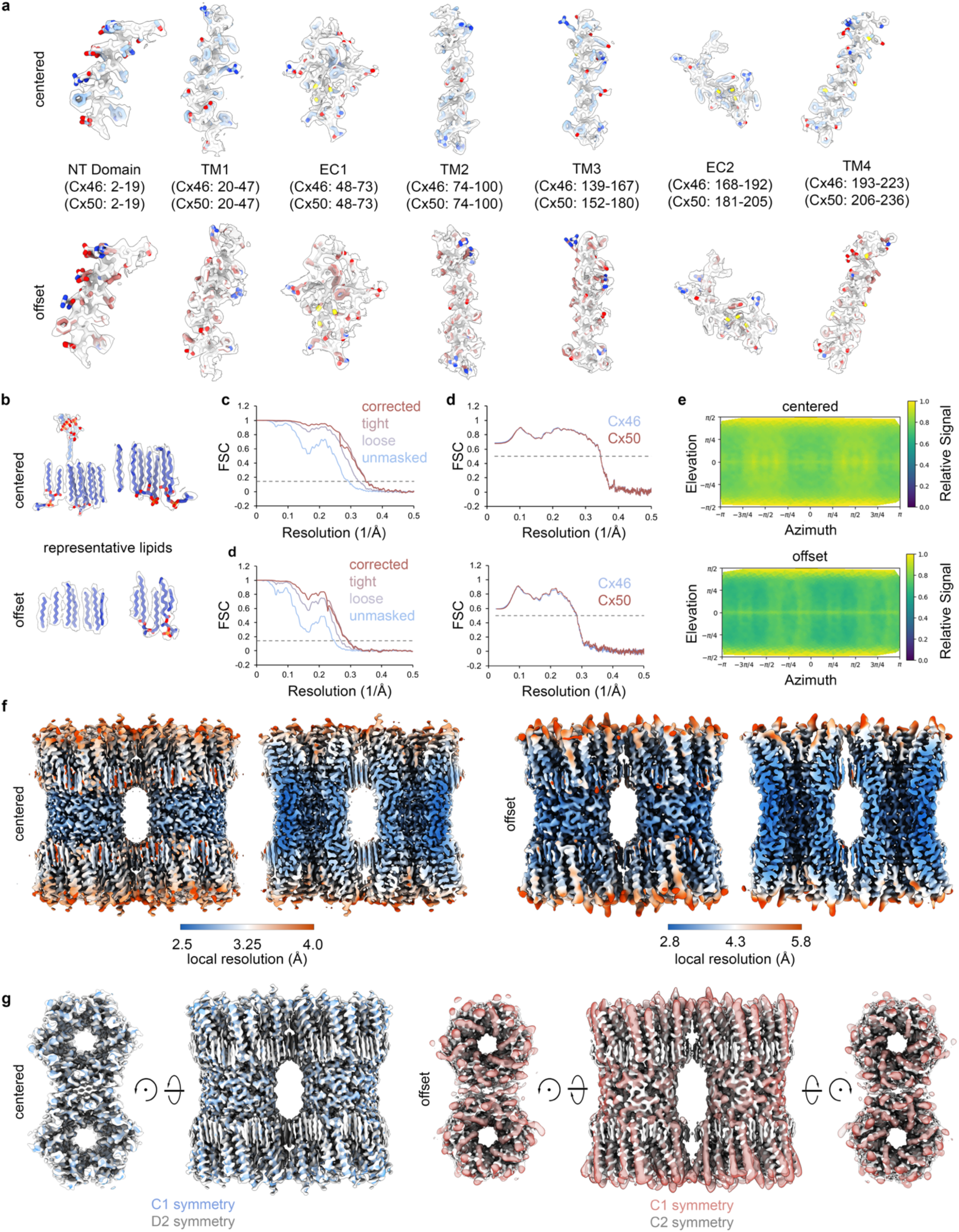
Map quality, validation, and local resolution of the centered and offset dual-channel reconstructions. **a,** Map-to-model fits for all structured regions of the centered and offset dual-channel reconstructions. Residue ranges for bovine Cx46 and Cx50 are indicated below each segment. Cx46 – white, Cx50 – blue (centered, top row) and red (offset, bottom row). **b,** Map-to-model fit of representative lipids for the centered and offset configurations. **c,** Gold-standard FSC curves (corrected, tight-mask, loose-mask, and unmasked), for the centered (top) and offset (bottom) reconstructions, with the 0.143 cutoff indicated by the gray dashed line. **d,** Unmasked map-to-model FSC curves for the Cx46 (blue) and Cx50 (red) models, for the centered (top) and offset (bottom) reconstructions. with the 0.5 cutoff indicated by the gray dashed line. **e,** Azimuth-elevation plots for the centered (top) and offset (bottom) reconstructions. Both particle sets show broad angular coverage. **f,** Cryo-EM maps colored by local resolution for the centered (left) and offset (right) reconstructions. In both assemblies, local resolution is highest in the central extracellular docking region and lower toward the cytoplasmic regions. **g,** Overlay of symmetrized and unsymmetrized reconstructions for each dual-channel configuration, showing that refinement with imposed symmetry does not alter overall channel architecture. For the centered configuration, the D2-symmetrized map is shown in white and the corresponding C1 map in blue. For the offset configuration, the C2-symmetrized map is shown in white and the corresponding C1 map in red.

**Supplementary Figure 6:**
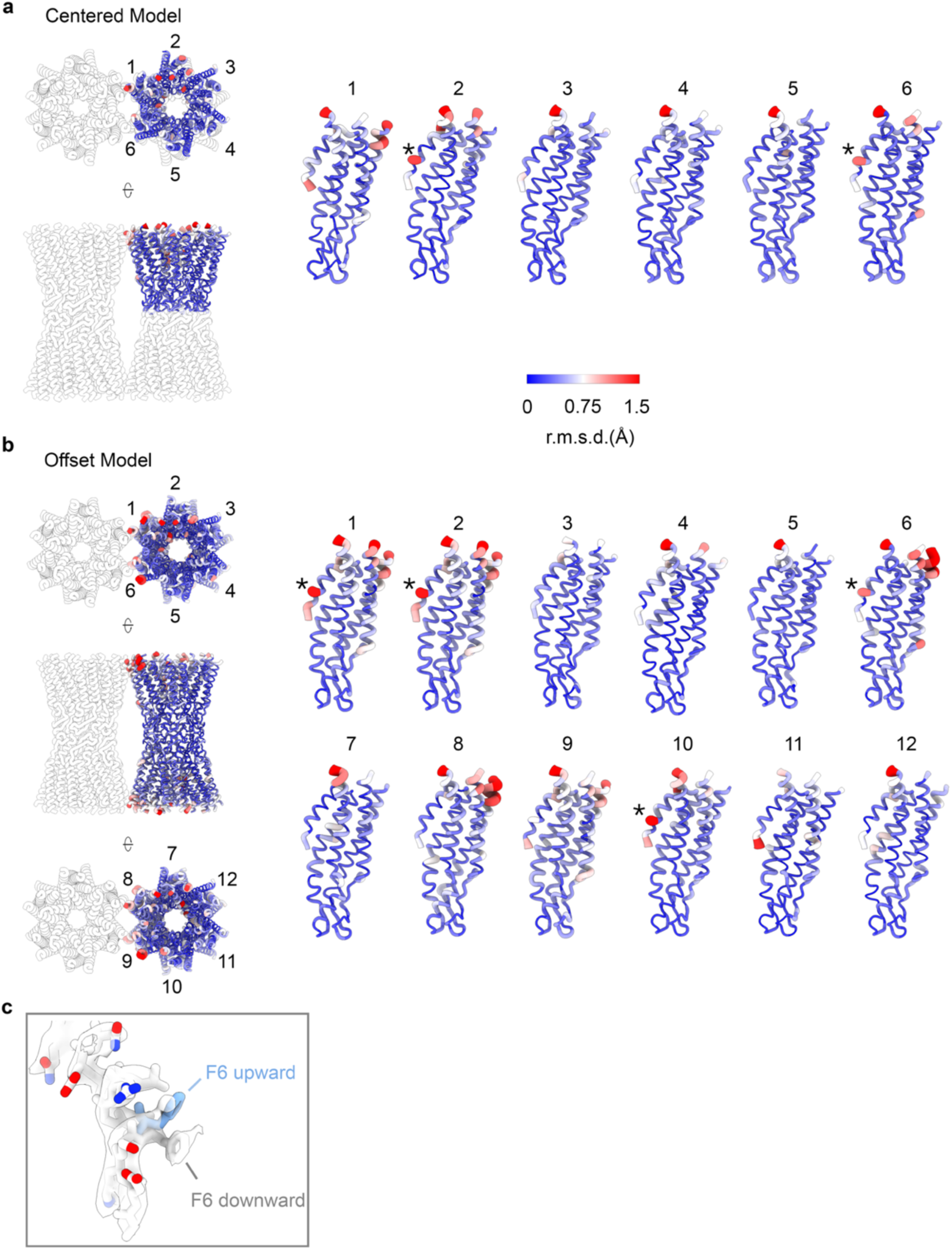
Open-state channel architecture is preserved in dual-channel conformations. **a,** Structural comparison of the centered dual-channel model to the 1.8 Å single-channel Cx46 model in the stabilized open-state conformation. The centered asymmetric unit (hemichannel) was aligned to the single-channel model, and Cα r.m.s.d. was mapped onto each of the six uniquely modeled subunits (1–6) in the hemichannel. **b,** Structural comparison of the offset dual-channel model to the 1.8 Å single-channel Cx46 model. The offset asymmetric unit (the complete intercellular channel) was aligned to the single-channel model, and Cα r.m.s.d. was mapped onto each of the twelve uniquely modeled subunits (1–12). In *(a)* and *(b)*, worm radius and color represent Cα r.m.s.d. (color scale shown). **c,** Example illustrating the primary localized difference contributing to higher r.m.s.d. in the NT region. In a subset of subunits, density supports an alternative rotamer position of F6 (“downward”; white) relative to the “upward” position observed in the single-channel model (blue) and most subunits in the dual-channel reconstructions.

**Supplementary Figure 7:**
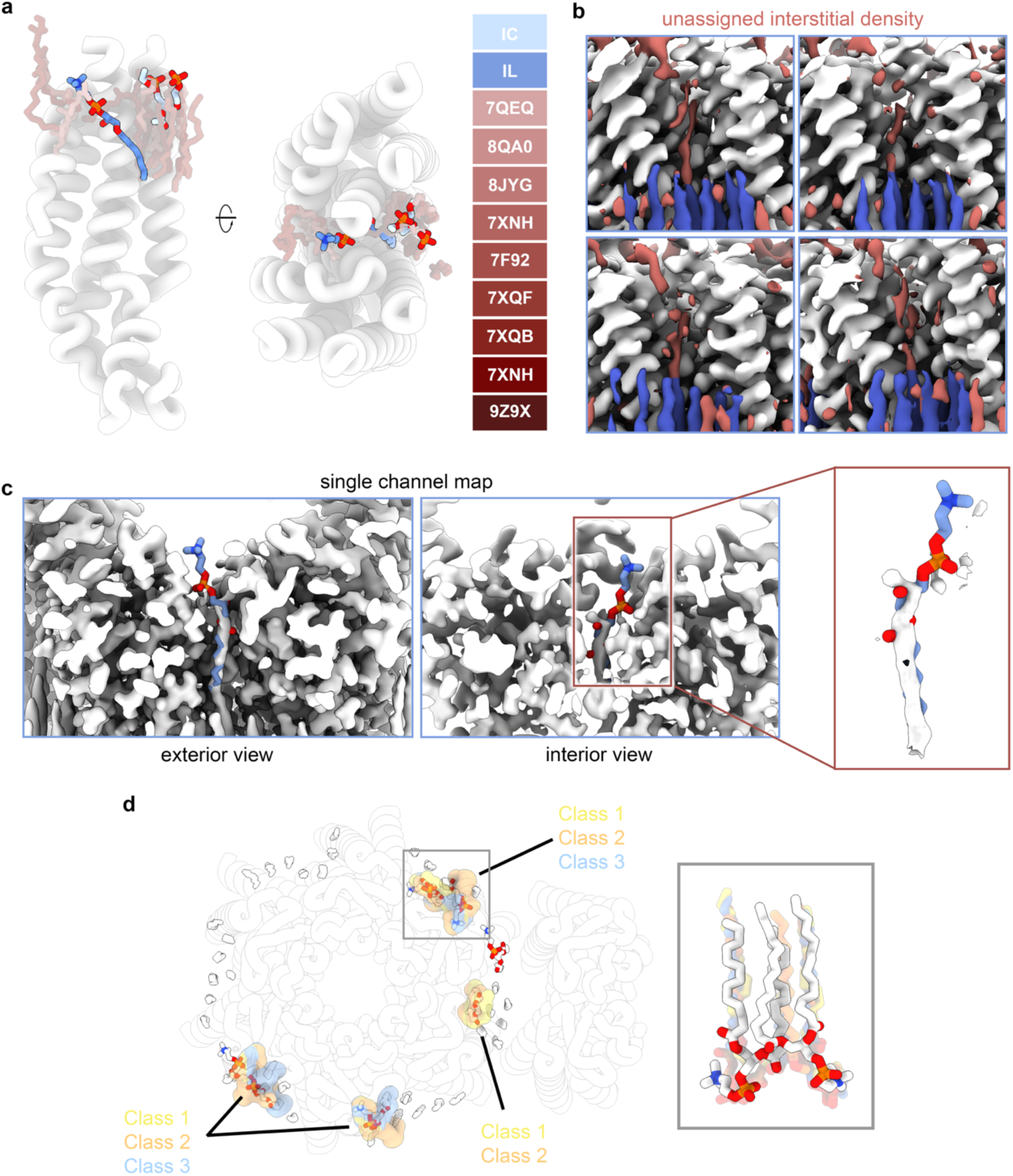
The dual-channel IL lipid occupies a unique position in the channel-channel interface. **a,** Comparison of lipid positions in the intracellular leaflet (IC) and interstitial lipid (IL) regions between this study and previously reported connexin structures. Lipids from this study are shown in blue, and representative lipids from previously published structures are shown in red (PDB IDs indicated). IC leaflet lipid headgroups occupy a commonly observed lipid-rich region, whereas the IL lipid resolved in the centered dual-channel interface occupies a distinct position. **b,** Unassigned density is present at IL-equivalent sites between other subunit interfaces in the centered dual-channel reconstruction but lacks sufficient connectivity and/or headgroup definition for confident modeling. Protein is shown in white, modeled lipid densities in blue, and unmodeled density in red. **c,** The IL headgroup is not resolved in the 1.8 Å single-channel reconstruction. Left,exterior view of the IL site. Middle, pore-facing view of the same region. Right, enlargement showing that residual density overlaps only with the acyl-chain portion of the IL site and does not support modeling of the IL lipid headgroup projecting toward the NT, as observed in the centered dual-channel interface. **d,** Extracellular leaflet (EC) lipid headgroups resolved in the centered dual-channel reconstruction align with previously reported EC lipid classes resolved for Cx46/50^16^. Headgroup-containing EC lipids from this study are shown as white sticks and overlaid with previously defined headgroup lipid classes from single-channel Cx46/50 reconstructions (Class 1, yellow; Class 2, orange; Class 3, blue; surfaces). *Inset*, representative EC lipids illustrating overlap of headgroups with prior class positions.

## SUPPLEMENTARY MOVIE LEGENDS

**Supplementary Movie 1. Morph between EMDB-22385 at 1.9 Å resolution and 1.8 Å single-channel map of Cx46/50.**

**Supplementary Movie 2. 3DVA separation of dual- and single-channel assemblies.**

**Supplementary Movie 3. 3DVA separation of centered and offset configurations.**

**Supplementary Movie 4. Morph between centered and offset atomic models.**

